# Tropomyosin 1-I/C co-ordinates kinesin-1 and dynein motors during *oskar* mRNA transport

**DOI:** 10.1101/2022.10.26.513919

**Authors:** Simone Heber, Mark A. McClintock, Bernd Simon, Janosch Hennig, Simon L. Bullock, Anne Ephrussi

## Abstract

Dynein and kinesin motors mediate long-range intracellular transport, translocating towards microtubule minus and plus ends, respectively. Cargoes often undergo bidirectional transport by binding to both motors simultaneously. However, it is not known how motor activities are coordinated in such circumstances. In *Drosophila*, sequential activities of the dynein-dynactin-BicD-Egalitarian (DDBE) complex and of kinesin-1 deliver *oskar* mRNA from nurse cells to the oocyte, and within the oocyte to the posterior pole. Here, through *in vitro* reconstitution, we show that Tm1-I/C, a Tropomyosin-1 isoform, links kinesin-1 in an inactive state to DDBE-associated *oskar* mRNA. NMR spectroscopy, small-angle X-ray scattering and structural modeling indicate that Tm1-I/C suppresses kinesin-1 activity by stabilizing its autoinhibited conformation, thus preventing a tug-of-war between the opposite polarity motors until kinesin-1 is activated in the oocyte. Our work reveals a novel strategy ensuring sequential activity of microtubule motors.

## Introduction

Motor protein driven transport along cytoskeletal tracks is important for subcellular localization of many cargoes, including organelles, vesicles and RNAs. Whereas myosin motors transport cargoes on actin filaments^1^, cytoplasmic dynein and kinesin motors are responsible for translocation along microtubules, driving minus-end-directed and plus-end-directed movements, respectively. The cargoes for these motors, however, often travel bidirectionally, requiring the coordinated activity of the two motor classes. For instance, in neurons, mRNAs^2,3^ and mitochondria^4^ are shuttled bidirectionally to ensure their constant supply to distal regions. Because dynein and kinesin motors are often simultaneously bound to the same cargo^5,6^, their activities must be tightly coordinated to allow for efficient transport rather than a futile tug-of-war.

Individual motors appear to be in an autoinhibited state by default, with activation driven by binding of adaptor proteins and cargoes. For instance, dynein is activated by binding of dynactin and one of a number of coiled-coil containing activating adaptors^7,8,9,10,11^. In the case of mRNA transport in *Drosophila*, dynein uses Bicaudal-D (BicD) as the activating adaptor, which is linked to double-stranded mRNA localization signals by the RNA-binding protein Egalitarian (Egl)^12,13^. Interaction of Egl and BicD with each other, as well as with the motor are enhanced by binding the mRNA signal, revealing a role for the mRNA in controlling motor activity^12,13,14,15^.

Kinesin-1 is a tetramer consisting of two Kinesin heavy chains (Khc) and two Kinesin light chains (Klc)^16^. Khc contains an N-terminal globular motor domain, a coiled-coil stalk domain and a C-terminal disordered tail domain (Figure 1a) and is autoinhibited by folding of the tail region of the heavy chain back on its motor domain. This interaction has been proposed to involve pivoting of the coiled-coil stalk at a hinge region, which allows docking of a conserved isoleucine-alanine-lysine (IAK) motif in the tail onto the motor domain, thereby locking the complex in a conformation that prevents processive movement^17,18,19,20^. The autoinhibited conformation is stabilized by Klc^21,22^, which binds to the Khc stalk domain^23,24^ and is important for recognition of a range of cargoes^25^.

**Figure 1:**
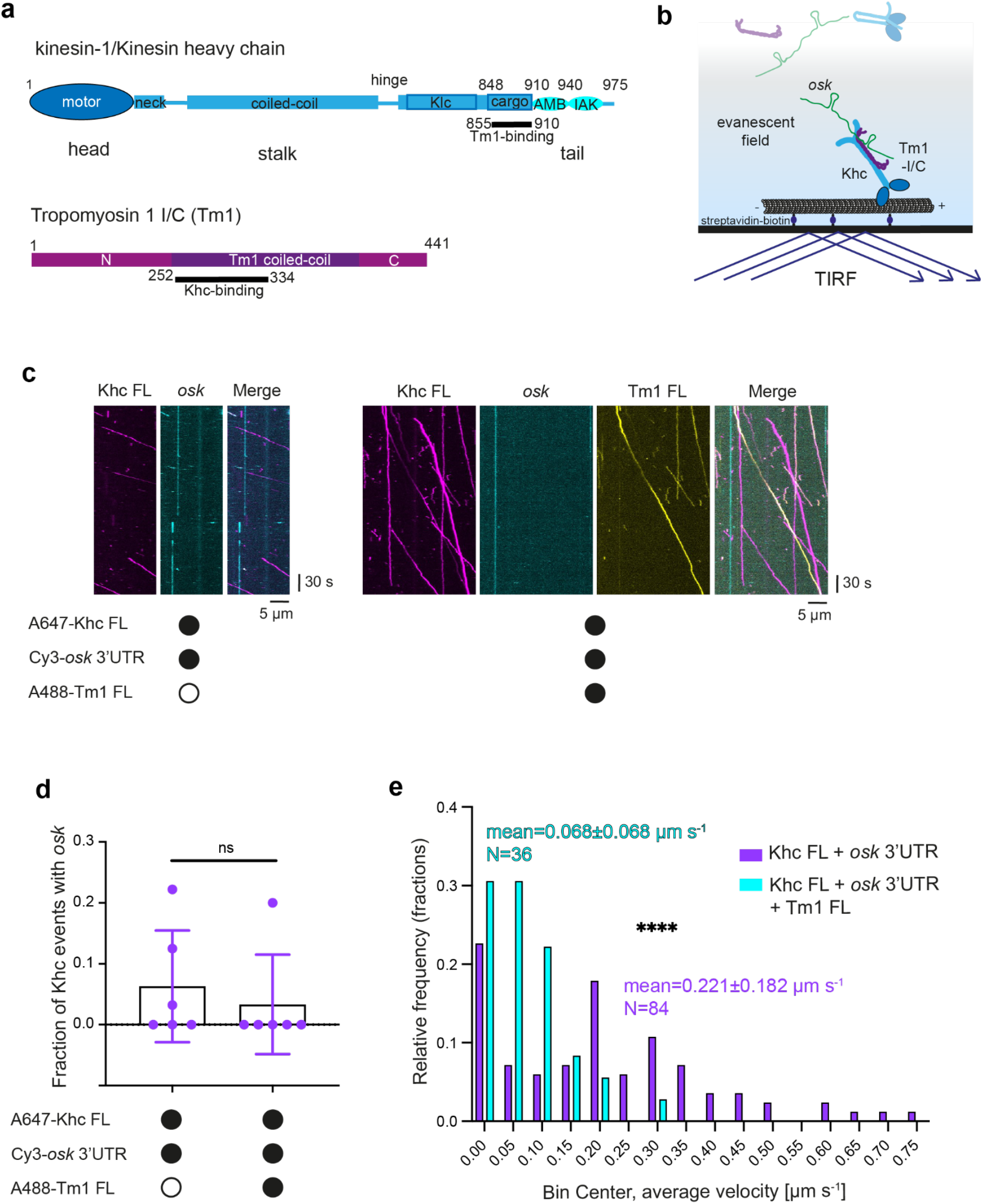
A TIRF microscopy-based *in vitro* motility assay reveals an inhibitory effect of Tm1 on Khc motility. **a** Scheme of domain architecture of *Drosophila* Kinesin heavy chain (Khc) and atypical Tropomyosin 1 isoform I/C (Tm1). **b** Scheme of *in vitro* motility assay. Preincubated Khc, Tm1 and RNA mixes were diluted to 6.25 nM of each component, and injected into an imaging chamber with pre-immobilised microtubules on a passivated glass surface. **c** Example kymographs showing processive movements of AlexaFluor647-labeled SNAP-Khc FL (labeling efficiency: 79.75 %), Cy3-labeled *osk* 3’UTR and AlexaFluor488-labeled Tm1 FL (labeling efficiency: 57 %). Microtubule plus and minus ends are oriented toward the right and left of each kymograph, respectively. **d** Fraction of Khc transporting *osk* 3’UTR in absence and presence of AlexaFluor488-labeled Tm1 in the assembly mix. Data points represent individual microtubules (84 and 36 complexes analyzed per condition). Statistical significance was evaluated with an unpaired t-test. ns = not significant. **e** Histograms of velocity distributions of Khc in absence or presence of Tm1. Statistical significance was evaluated with an unpaired t-test. ****p<0.0001.

However, the mechanisms of regulation in dual motor transport remain elusive, with only a small number of adaptor proteins that coordinate both motors having been reported. For example, the mitochondrial transport adaptor TRAK2 forms a complex with both kinesin-1 and dynein-dynactin and links these two motors functionally^26,27^. UNC-16, the *C. elegans* homolog of the lysosome adaptor JIP3/Syd, was also shown to link kinesin-1 and dynein, allowing for plus-end-directed transport of dynein by kinesin-1^28,29^. Similarly, the dynein adaptor Hook3 can activate and link KIF1C (a kinesin-3 motor) and dynein^30,31^.

*oskar* (*osk*) mRNA localization in the *Drosophila* egg chamber is a paradigm for the study of RNA transport and motor regulation. Motor-based delivery of *osk* to the posterior pole of the developing oocyte drives abdominal patterning and germline formation in the embryo^32^. Trafficking of *osk* mRNA to this site is mediated by the successive activities of the dynein^33^ and kinesin-1^34.35^ motors, and requires motor switching. In the germline syncytium, the oocyte is transcriptionally quiescent and relies on mRNAs provided by the 15 interconnected nurse cells for its development^36^. At this stage, microtubule minus ends are nucleated in the oocyte and extend into the nurse cells. Upon transcription in the nurse cells, *osk* mRNA is transported along the polarized microtubule cytoskeleton into the oocyte by dynein in complex with dynactin, BicD and Egl^12,33,37^. During mid-oogenesis, polarity of the microtubule cytoskeleton changes dramatically, with microtubule plus ends becoming enriched at the oocyte posterior. At this point, Khc translocates *osk* to the posterior pole^38^. Khc was previously shown to function in *Drosophila* oocytes largely independently of Klc^39,40^, raising the question of how Khc is linked to *osk* and how motor activity is regulated.

Kinesin-mediated transport of *osk* RNA requires the unique I/C isoform of tropomyosin-1, Tm1-I/C (hereafter referred to as Tm1)^39,40,41,42^. Tm1 binds to a non-canonical but conserved cargo binding region in the Khc tail and stabilizes the interaction of the motor with RNA, suggesting a function as an adaptor^43^. Both Khc and Tm1 are already loaded onto the *osk* ribonucleoprotein particle (RNP) shortly after nuclear export in the nurse cells, although the motor only appears to become active in the oocyte during mid-oogenesis^42^. Similarly, dynein remains associated with *osk* RNPs during Khc-mediated transport within the oocyte, but is inactivated by displacement of Egl by the Staufen (Stau) protein^44,45^.

How the two motors are linked simultaneously to *osk* RNPs, and how cargo-bound Khc is inhibited during dynein-mediated transport into the oocyte has been unclear. Here, we show that Tm1 inhibits Khc activity by stabilizing its autoinhibited conformation through a novel mechanism involving the regulatory tail domain. In addition, Tm1 links Khc to the dynein-transported *osk* RNP, thereby allowing co-transport of inactive Khc on *osk* RNA by dynein. *In vivo*, such a mechanism would avoid a tug-of-war between the two motors during the dynein-dependent delivery of *osk* RNPs to the oocyte whilst ensuring that Khc is available on these structures when it is time for this motor to move the RNPs to the posterior of the oocyte. With its cargo-binding and motor regulatory functions, we propose that Tm1 functions as a non-canonical light chain for kinesin-1 in the *Drosophila* oocyte.

## Results

### Tm1 can transport RNA with Khc *in vitro* but impairs Khc motility

In order to test the ability of Khc, Tm1 and the Khc-Tm1 complex to transport RNA, we performed total internal reflection fluorescence (TIRF) microscopy-based *in vitro* motility assays^15^ using fluorescently labeled versions of recombinant, full-length Khc (Khc FL), full-length Tm1 (Tm1 FL) (Figure 1a, b) and *in vitro* transcribed *osk* 3’UTR RNA. At concentrations that allowed single-particle tracking (6.25 nM of each component), we observed transport of *osk* by Khc along microtubules in both the presence and absence of Tm1, albeit at low frequency (Figure 1c). In the absence of Tm1, 4.8 % (4/84) of Khc transport events co-localized with the *osk* 3’UTR signal. This value did not change significantly in the presence of Tm1 (2.8 %, 1/36; Figure 1d), despite 55 % (20/36) of Khc signals co-localizing with Tm1 signals. Correcting for the efficiency of labeling of Tm1 with the fluorophore (57 %) indicated that ∼95 % of microtubule-bound Khc molecules were bound to Tm1. The high occupancy of Khc with Tm1 in these experiments is consistent with the strong binding between their interacting domains observed previously^43^. Mass photometry confirmed high occupancy of Khc with Tm1 at nanomolar concentrations (Figure S1), as well as the proposed 2:1 stoichiometry of the Khc-Tm1 complex^43^. We conclude from these experiments that Khc can associate with *osk* RNA but that the Khc-Tm1 complex does not have substantially higher affinity for the transcript in our assay conditions.

We next examined the effect of Tm1 on Khc motility. Without Tm1 present in the assay, microtubule-associated Khc complexes moved with a mean velocity of 0.221 µm s^-1^. In the presence of Tm1, the mean velocity of Khc runs reduced to 0.068 µm s^-1^ (Figure 1e). These data reveal an inhibitory effect of Tm1 on Khc motility.

In addition to the 3’UTR, an RNA structure in the *osk* coding sequence, which is called the spliced *oskar* localization element (SOLE), is important for Khc-mediated transport of *osk in vivo*^46,47^. We tested if an RNA including the SOLE could stimulate motility of Khc in the presence of Tm1 by replacing the *osk* 3’UTR in our assay with a 135-nt RNA (*osk* min) comprising the SOLE and the oocyte entry signal of *osk* (Figure S2a), which resides in the 3’UTR and promotes dynein-mediated transport^48^. The inhibitory effect of Tm1 on Khc motility was also observed in the presence of *osk* min (Figure S2b-f). Thus, Tm1 inhibits motility of Khc in both the presence and absence of the SOLE element.

### Tm1 inhibits Khc motility via the Khc regulatory tail

In order to elucidate the molecular mechanism behind Tm1-mediated inhibition of Khc, we set out to identify the region of Khc that is involved in this process. Regulatory functions have been attributed to an auxiliary, ATP-independent microtubule-binding domain (AMB) and the autoinhibitory IAK motif within the Khc tail (Figure 1a)^19,49,50^. Whereas the AMB allows for microtubule-sliding by Khc^51^, the IAK motif inhibits Khc’s ATPase activity^17,19^, and thus processive motility, by binding to the Khc motor domain and locking the two Khc heads in a conformation that prevents neck-linker undocking and ADP-release^18^. We tested whether the regulatory tail is responsible for Tm1-mediated inhibition of Khc by assessing the effect of Tm1 on the motility of a tailless Khc truncation (Khc910; Figure 2a). As expected of a Khc without its regulatory tail, Khc910 was hyperactive in the *in vitro* motility assay, exhibiting more processive runs (Figure 2c and Table S1), an increased likelihood of processive movement upon binding microtubules (Figure 2d), and faster motility (Figure 2e and Table S1) than the full-length motor complex. However, the presence of Tm1 did not inhibit Khc910 motility (Figure 2c-e and Table S1). The same effects were observed when experiments were performed with *osk* min instead of *osk* 3’UTR (Figure S2c-f). These observations demonstrate that the regulatory tail of Khc is required for Tm1-mediated inhibition of motility.

**Figure 2:**
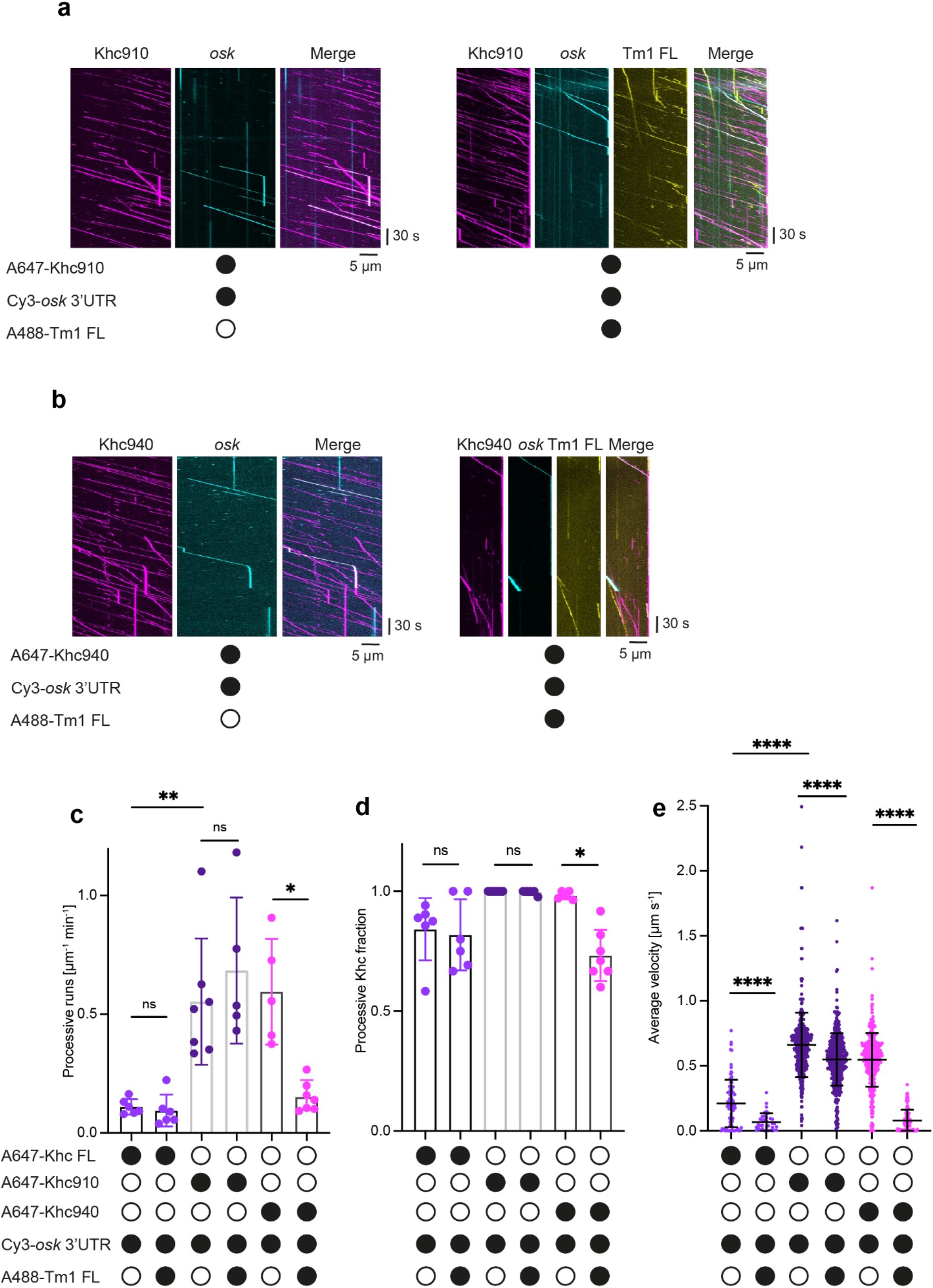
Tm1 inhibits Khc motility *in vitro* via Khc’s regulatory tail. **a, b** Example kymographs showing processive movements of AlexaFluor647-labeled SNAP-Khc910 (a) and Khc940 (b) ± AlexaFluor488-labeled Tm1 FL. Microtubule plus and minus ends are oriented toward the right and left of each kymograph, respectively. **c, d** Number of processive events of Khc FL, Khc940 and Khc910 (c) and fraction of microtubule-bound Khc complexes that are processive ± Tm1 FL in the assembly mix (d). Data points show values for individual microtubules (between 36 and 438 complexes analyzed per condition). Error bars: SD. Statistical significance was evaluated with an unpaired t-test. ns = not significant, **p<0.01, *p<0.05. Labeling efficiencies for Khc FL, Khc910 and Khc940 were 79.8 %, 54 % and 78 %, respectively, so that total run numbers are underestimated. **e** Average velocities of Khc FL and Khc910 ± Tm1 FL in the assembly mix. Data points represent individual runs. Statistical significance was evaluated with an unpaired t-test. ****p<0.0001.

To assess the relevance of the autoinhibitory IAK motif in the Khc tail for Tm1-mediated inhibition, we deleted this feature whilst leaving the AMB intact. To this end, we produced Khc940, which is truncated C-terminal to the AMB domain. Khc940 was hyperactive (in terms of the number of processive runs, fraction of Khc binding events that resulted in processive motility, and velocity) to a degree comparable to that of Khc910 (Figure 2b-e). However, despite the fact that a smaller fraction of Khc940 co-localized with Tm1 as compared with Khc FL and Khc910 (Figure S3), Khc940 was sensitive to inhibition by Tm1, as the number of runs, fraction of processive Khc, and Khc velocities were reduced in its presence (Figure 2b-e and Table S1). We conclude that Khc inhibition by Tm1 does not depend on the IAK motif and therefore involves another region of the Khc tail.

### Khc inhibition by Tm1 is mediated by the Khc AMB domain

To test if the AMB domain in the Khc tail is required for inhibition by Tm1, we produced a Khc mutant in which only the AMB domain (aa 911-938) was deleted (KhcΔAMB) and tested its motility with and without Tm1 (Figure 3a). KhcΔAMB did not have an increased frequency or likelihood of processive movement or higher velocity compared to Khc FL (Figure 3b-d and Table S2), indicating that autoinhibition is not disrupted in this protein. Strikingly, the frequency, likelihood of motility, and velocity of processive KhcΔAMB movements were not decreased by Tm1 (Figure 3a-d and Table S2), despite the proportion of complexes associated with Tm1 not differing significantly from what was observed for Khc FL or Khc910 (Figure S4). Taken together, these results reveal that the AMB domain of Khc mediates inhibition by Tm1 but is not necessary for Tm1 binding.

**Figure 3:**
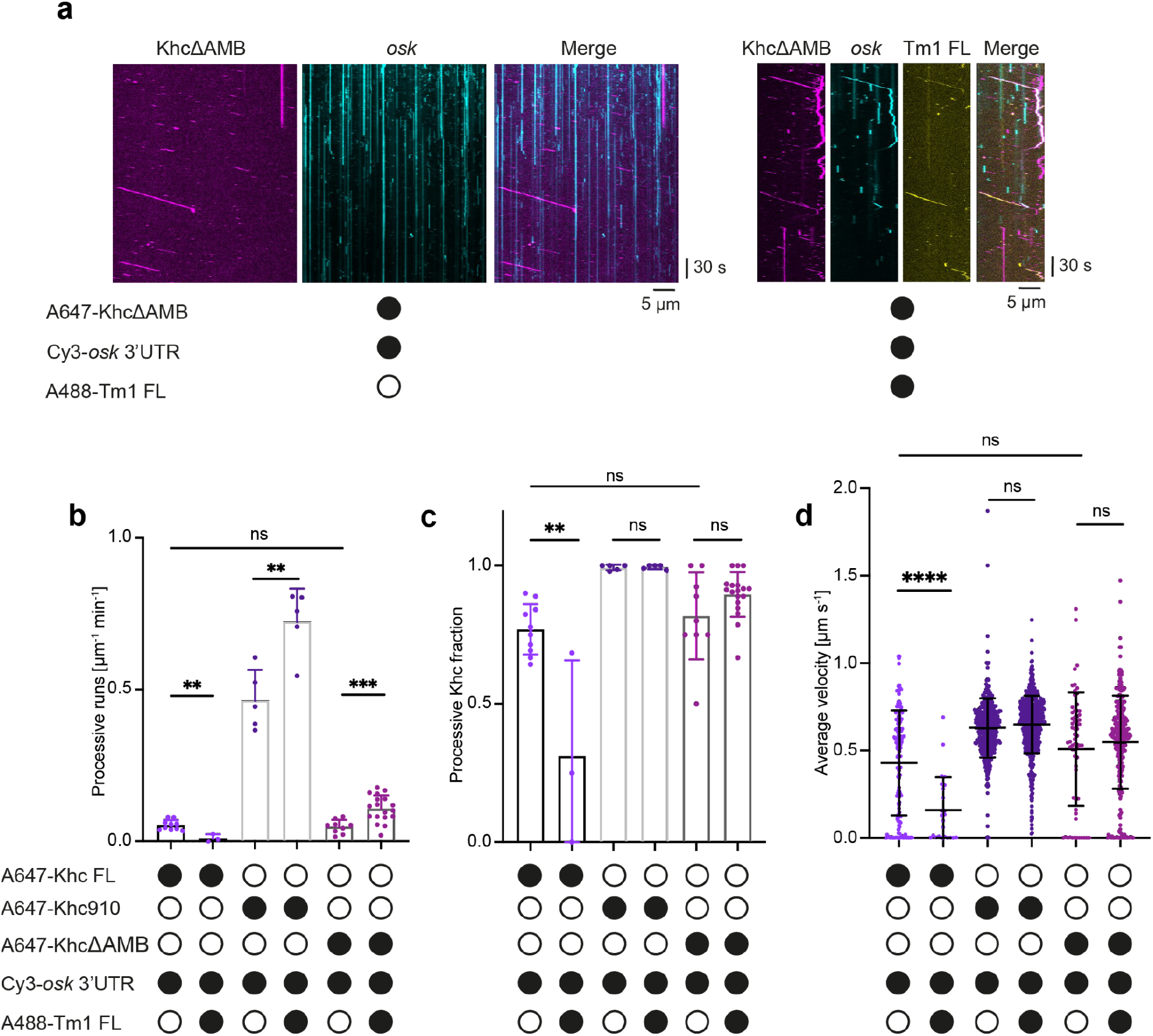
Khc AMB domain mediates Tm1 inhibition of Khc. **a** Example kymographs showing processive movements of AlexaFluor647-labeled SNAP-KhcΔAMB ± AlexaFluor488-labeled Tm1 FL in the assembly mix. Microtubule plus and minus ends are oriented toward the right and left of each kymograph, respectively. **b, c** Processive events of Khc FL, Khc910 and KhcΔAMB ± Tm1 FL (b) and fraction of processive Khc events (c). Data points show values for individual microtubules (between 24 and 595 complexes analyzed per condition). Error bars: SD. Statistical significance was evaluated with an unpaired t-test. ****p<0.0001, ***p<0.001, **p<0.01. Labeling efficiencies for Khc FL, Khc910 and KhcΔAMB were 79.8 %, 54 % and 78.8 %, respectively, so that total run numbers are underestimated. **d** Average velocities of Khc FL, Khc910 and KhcΔAMB ± Tm1 FL. Data points represent individual runs. Statistical significance was evaluated with an unpaired t-test. ****p<0.0001.

### The N-terminal domain of Tm1 is required for its inhibitory function

We next investigated which features of Tm1 mediate Khc inhibition. We previously showed that a truncated Tm1 protein (1-335 of the 441-amino-acid full-length protein) comprising the disordered N-terminal domain and the minimal Khc-binding region of the coiled-coil domain can suppress *osk* RNA mislocalization in *Drosophila* oocytes that lack the endogenous Tm1 protein^43^. We therefore compared the effect of Tm1 1-335 to Tm1 FL in the motility assay (Figure 4a) and found that both proteins inhibit Khc motility to a similar extent (Figure 4b-d and Table S3). We also observed that the fraction of Tm1-bound Khc did not differ significantly between Tm1 1-335 and Tm1 FL (Figure S5). These data show that the N-terminal region of Tm1 is sufficient to inhibit Khc.

**Figure 4:**
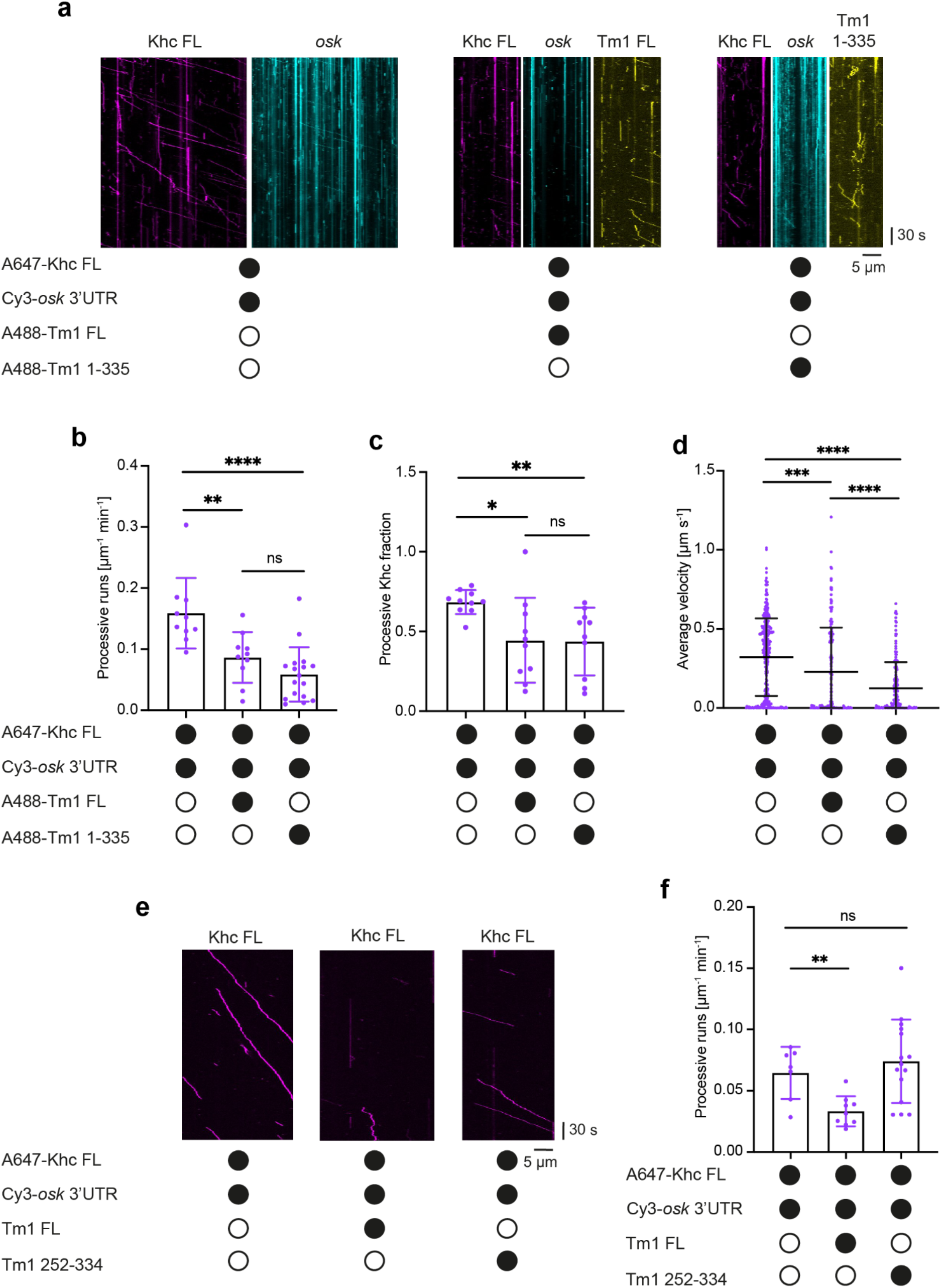
The Tm1 N-terminal region is required for Khc inhibition. **a** Example kymographs showing processive movements of AlexaFluor647-labeled SNAP-Khc FL ± AlexaFluor488-labeled Tm1 FL or Tm1 1-335 in the assembly mix. **b, c** Processive events of Khc FL ± Tm1 FL or Tm1 1-335 in the assembly mix (b) and fraction of processive Khc events (c). Data points show values for individual microtubules (between 44 and 335 complexes analyzed per condition). Error bars: SD. Statistical significance was evaluated with an unpaired t-test. ****p<0.0001, ***p<0.001. **d** Average velocities of Khc FL ± Tm1 FL or Tm1 1-335 in the assembly mix. Data points represent individual runs. Statistical significance was evaluated with an unpaired t-test. ****p<0.0001. **e** Example kymographs showing processive movements of AlexaFluor647-labeled SNAP-Khc FL ± Tm1 FL or Tm1 252-334. **f** Processive events of Khc FL ± Tm1 FL or Tm1 252-334 in the assembly mix. Error bars: SD. Statistical significance was evaluated with an unpaired t-test with Welch’s correction. **p<0.01. Microtubule plus and minus ends are oriented toward the right and left of each kymograph, respectively (a and e).

To address whether the Khc-binding domain of Tm1 (aa 252-334) is also sufficient to produce an inhibitory effect we performed motility assays with this protein. We compared the effects of untagged, unlabeled Tm1 FL and Tm1 252-334 (Figure 4e) on Khc motility. Whereas Tm1 FL again decreased the number of processive runs of Khc, we observed no negative effect of Tm1 252-334 on motility (Figure 4e, f, Figure S5d-f and Table S3). These results suggest that inhibition of Khc requires the unstructured, N-terminal domain of Tm1.

### Tm1 binding affects the disordered region of the regulatory Khc tail

Next, we set out to better understand the molecular mechanism of Khc inhibition by Tm1. Because the inhibitory effect of Tm1 is mediated by the auxiliary microtubule-binding region (AMB) within the Khc tail, we reasoned that Tm1 could alter microtubule binding by the tail domain, perhaps by providing an additional interaction with microtubules. In order to test this notion, we performed microtubule-binding assays^52^ with recombinant Tm1 proteins. Tm1 was incubated with polymerized microtubules before pelleting of microtubules, together with bound protein, by ultracentrifugation. As a positive control, we confirmed that both Khc FL and Khc tail partially partitioned into the microtubule-containing pellet fraction (Figure S6a). A small fraction of Tm1 1-335 also partitioned into the pellet, indicating that the N-terminal or coiled-coil domain of Tm1 can interact with microtubules. However, we did not observe increased partitioning of the Khc tail (Khc 855-975) into the microtubule fraction in the presence of Tm1 1-335 (Figure S6b). This observation indicates that association with Tm1 does not alter the degree of binding of the Khc tail to microtubules.

In the crystal structure of the Khc tail bound to Tm1, the Khc coiled coil extends into the AMB domain (last visible aa: 916)^43^. Attempts to crystallize the same Khc construct alone were unsuccessful, raising the possibility that the isolated motor protein is more flexible. It is conceivable that the extended coiled coil of Khc is induced upon Tm1 binding, which could reduce flexibility of the tail, potentially locking it in a conformation that impairs Khc activity. To test this hypothesis, we probed the structure of the Khc alternative cargo-binding domain and tail (Khc 855-975) in solution by NMR spectroscopy. The ^1^H,^15^N-HSQC NMR spectrum of the Khc tail showed 34 resonances, each corresponding to a backbone N-H correlation (Figure 5a). Backbone assignment experiments and resulting secondary chemical shifts revealed that the visible peaks are disordered and correspond to aa 937-975, the very C-terminal part of the Khc tail that contains the auto-inhibitory IAK motif. For all tested coiled-coil constructs of Khc, resonances for aa 855-936 were broadened beyond detection due to short transversal relaxation times. The unfavorable relaxation behavior of H-N resonances in coiled-coil structures is caused by slow rotational tumbling of the axis perpendicular to the helix orientation which is, in the case of Khc-Tm1, potentially further slowed down by transient intermolecular oligomerization. The absence of resonances for aa 855-936 is thus consistent with this entire part of the protein forming an extended coiled-coil structure, even in the absence of Tm1. Our data therefore suggest that the AMB domain is part of the structured coiled coil even in absence of Tm1.

**Figure 5:**
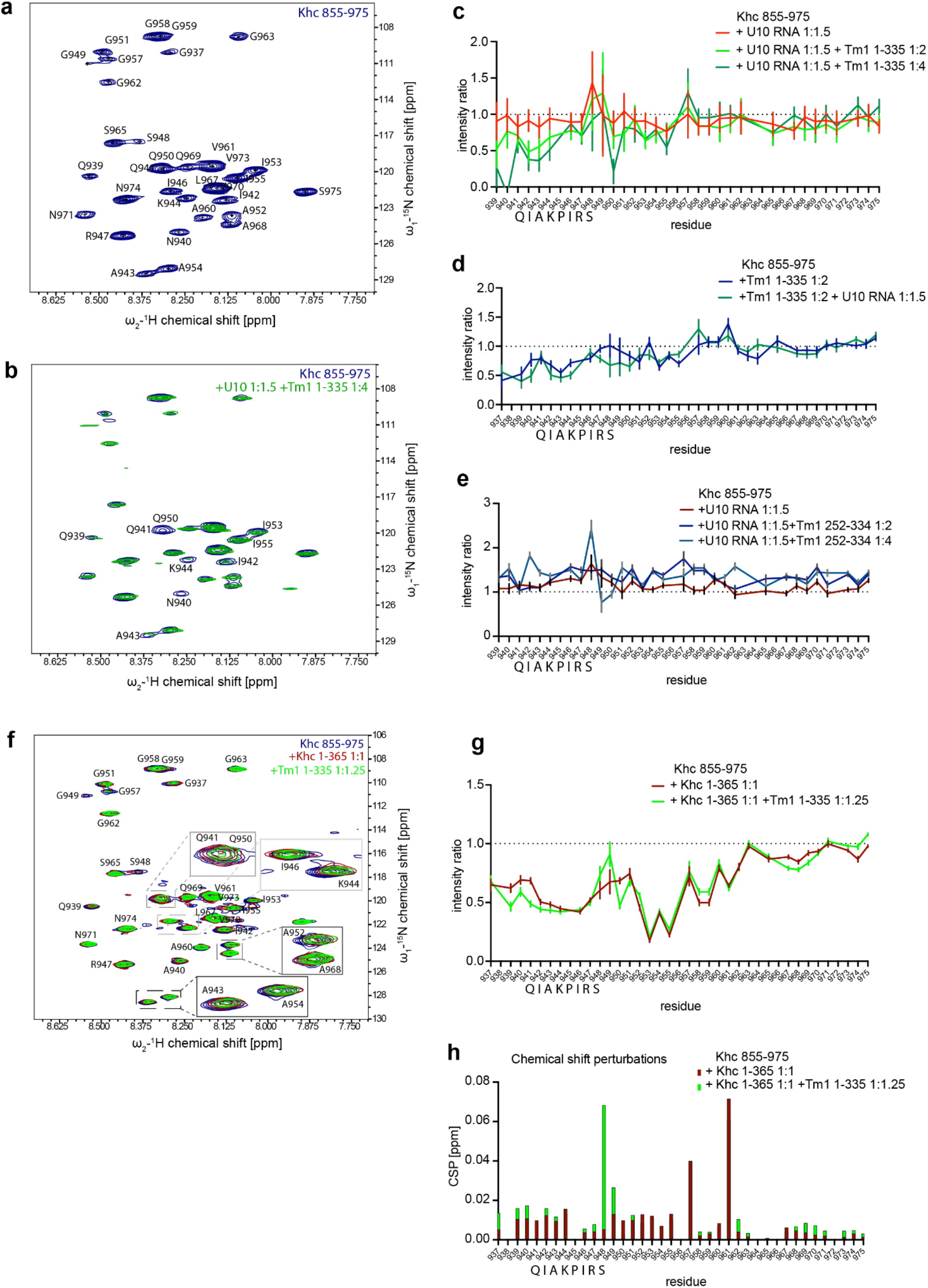
Tm1 binding affects the free and motor-bound regulatory tail of Khc. **a** ^1^H,^15^N-HSQC spectrum of the Khc tail construct Khc 855-975 with backbone assignments indicated. **b, c** ^1^H,^15^N-HSQC titration experiment with U10 RNA and Tm1 1-335. Overlaid spectra (b) and plotted intensity ratios of individual residues. RNA was added before Tm1. **d** Intensity ratios of a titration experiment with Tm1 1-335 and U10 RNA in which Tm1 was added before RNA. **e** Intensity ratios of a titration experiment with U10 RNA and Tm1 252-334 in which RNA was added before Tm1. **f - h** ^1^H,^15^N-HSQC titration experiment of the Khc tail Khc 855-975 with the motor domain Khc 1-365 and Tm1 1-335. Overlaid spectra (f), plotted intensity ratios (g) and chemical shift perturbation plot (h) of individual residues. Khc 1-365 was added before Tm1 1-335.

To further investigate the effect of Tm1-binding on the disordered tail, we performed NMR titration experiments to detect any effects of Tm1 binding on the tail region. Upon titration of increasing amounts of Tm1, there were intensity decreases for several resonances in ^1^H,^15^N-HSQC spectra of Khc 855-975 (Figure 5b, c). The affected residues correspond to the IAK motif and its flanking regions. This effect could be observed both in the absence and presence of a U10 RNA (Figure 5c, d), and depended on the N-terminal domain of Tm1 because the Tm1 coiled-coil (aa 252-334) construct did not affect the spectra (Figure 5e). These findings are in agreement with our earlier findings that Tm1 inhibits Khc motility in the absence of RNA and that the N-terminal region of Tm1 is required for its inhibitory effect. These intensity decreases for the IAK motif and its proximal residues indicate an induced conformational change or a second, transient interaction of the Tm1 N-terminal domain with this region.

### Tm1 does not affect the interaction of the Khc tail with the isolated Khc motor domain

The above results led us to hypothesize that Tm1, upon binding to the coiled coil of Khc that includes the AMB domain, influences the interaction of the Khc tail and motor domains. To test this notion, we performed NMR titration experiments in which the Khc motor domain (aa Khc 1-365) was added to the tail domain prior to Tm1 1-335 titration. Titration of Khc 1-365 to Khc 855-975 caused intensity losses for several resonances. As expected, the resonances corresponding to the IAK motif (aa 941-948), as well as its flanking residues (aa 936-962), were affected (Figure 5f, g). Some resonances in this region also showed chemical shift perturbations (Figure 5f, h). Taken together, these data confirm an interaction between Khc 855-975 (tail) and Khc 1-365 (motor). Additional titration of Tm1 1-335 to this Khc tail-motor complex caused further intensity decreases of the same resonances (Figure 5f, g), showing that Tm1 1-335 can bind to the Khc tail when it is associated with the motor domains. We conclude that binding of the Khc tail to Tm1 and the Khc motor are not mutually exclusive.

We therefore tested whether Tm1 can directly affect the interaction of the Khc motor and tail domains in GST-pulldown assays (Figure S7). When used as bait, the GST-tagged Khc tail domain (Khc 855-975) simultaneously captured the Khc motor domain (Khc 1-365) and Tm1 1-335, consistent with our NMR-based evidence that formation of a trimeric complex is possible. However, the fractions of Khc 1-365 and Tm1 1-335 pulled down by GST-Khc 855-975 were low, indicating a rather weak or transient interaction. Upon titration of Tm1 1-335, the amount of motor domain associating with the tail domain was not affected. Thus, we conclude that Tm1 does not alter the ability of the isolated Khc motor domain to interact with the isolated Khc tail domain.

### Tm1 alters the auto-inhibited conformation of Khc

We next reasoned that Tm1 binding might stabilize the autoinhibited conformation of the full-length Khc by favoring additional interactions within the coiled-coil stalk domain, likely including the AMB domain. In order to obtain information about Khc’s predominant conformation in its free and Tm1-bound states, we measured small-angle X-ray scattering (SAXS) of Khc and the Khc-Tm1 complex (Figure 6a). A dimensionless Kratky plot^53,54^ of both scattering profiles (Figure 6b) shows that Khc and Khc-Tm1 are extended molecules with flexible elements. To identify structures that can explain the scattering curves of Khc and Khc-Tm1, we generated a large pool of Khc and Khc-Tm1 models by connecting coordinates of the structured parts that were generated by AlphaFold2^55,56^. Based on AlphaFold2 predictions, the Khc stalk contains four coiled-coil regions (cc 1-4) connected by flexible hinge regions (hinges 2-4). The stalk is connected to the motor domain’s neck coiled coil by another flexible region (hinge 1) (Figure S8a) We subsequently randomized the connecting linker and tail regions to create a large pool of possible structures. In a second step, the C-terminal tail of one chain was pulled into the binding pocket of the motor domain resulting in a second pool of structures with random orientation of the stalk region and the IAK peptide bound to the motor domain dimer. Both pools of structures were sorted by the quality of the fit to the SAXS data.

**Figure 6:**
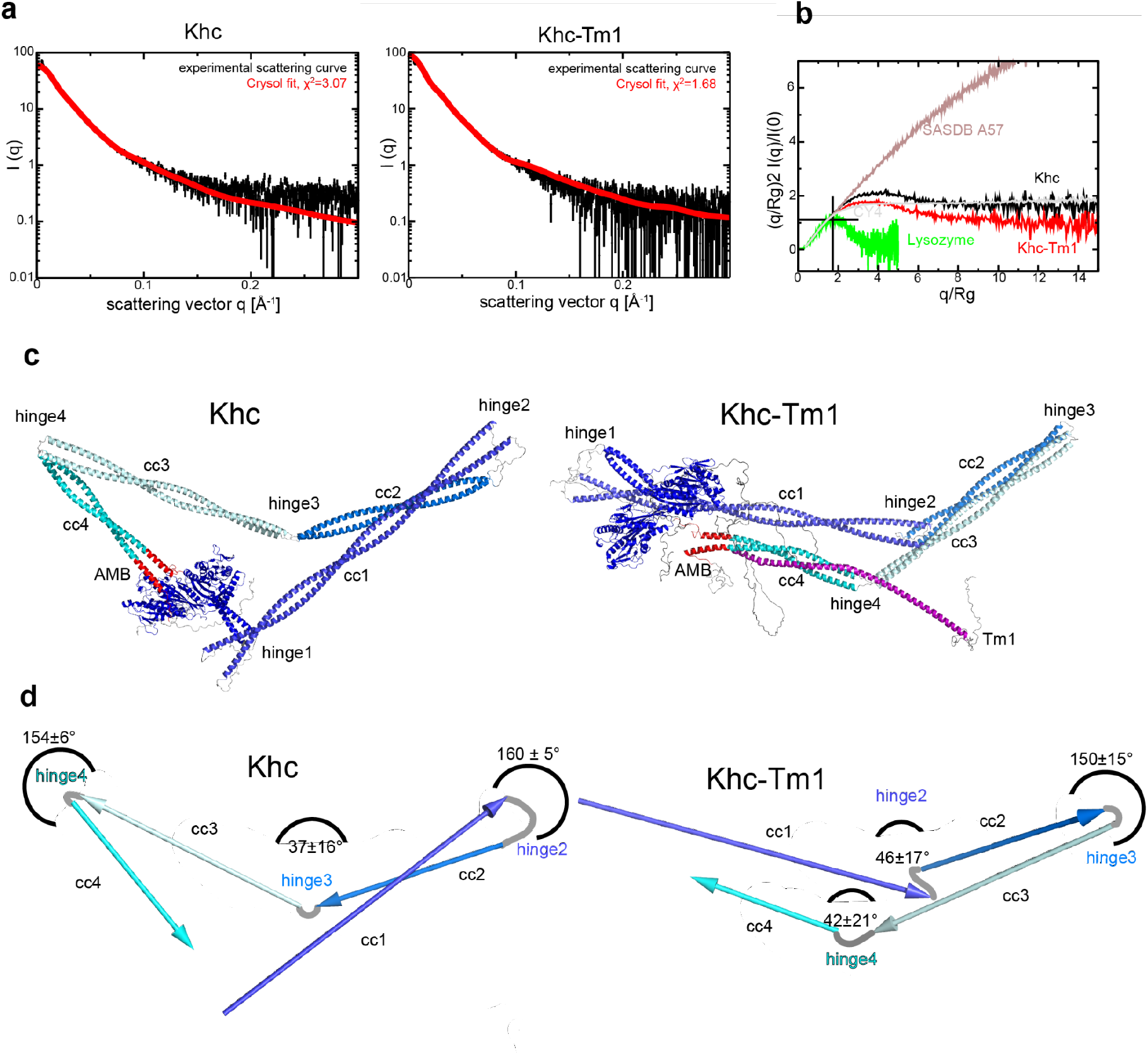
Khc conformation is altered upon Tm1 binding. **a** Experimental SAXS curves overlaid with theoretical SAXS curves of best Crysol fits. **b** Dimensionless Kratky plot of scattering profiles of Khc and the Khc-Tm1 complex compared to a globular protein (lysozyme), a disordered protein with short secondary structure elements (SASBDB^57^ CY4) and a fully disordered extended protein (SASBDB^57^ A57) to illustrate the differences between Khc and Khc-Tm1 and the contributions of ordered and disordered regions. **c** Best fits for Khc and for Khc-Tm1. Khc models are shown in shades of blue with the AMB domains marked in red, Tm1 is shown in pink. cc 1-4 denote the four coiled-coil regions of the Khc stalk domain. **d** Schemes showing the orientations of Khc’s stalk coiled coils in the structures with best Crysol fits. Angles between coiled coils are averages from the five best-fitting models.

For both Khc and the Khc-Tm1 complex, the structures with the lowest χ^2^ values (i.e. best quality fits) all exhibited compact conformations with considerably lower than average values for the radius of gyration (*Rg*). Since the autoinhibited conformations were more compact than the initial pool with randomized linker conformations, there were on average more structures with low χ^2^ values in the pools of structures with the IAK motif bound to the motor domains (Figure S8b). Inspection of the structural models that best fit the scattering curves indicated that the arrangement of the coiled-coils is different for Khc and Khc-Tm1. Looking at the angles (Figure 6c-d) of consecutive coiled-coil regions of Khc models, we observed that the conformation of the hinge regions (hinge 2: aa 586-603, hinge 3: aa 707-710 and hinge 4: aa 839-843, Figure S8a) leads to bent structures of the coiled-coil regions cc1-cc2 and cc3-cc4 and elongated cc2-cc3 orientation, resulting in a triangular shape of the molecule (Figure 6c, Figure S8d). In this conformation, the N- and C-terminal parts of the stalk are separated, and the AMB domain, which mediates inhibition by Tm1, is not in proximity to other stalk regions (Figure 6c, Figure S8d-e). For Khc-Tm1, the stalk folds at hinge 3 (Figure 6c, Figure S8d), bringing N- and C-terminal halves of the stalk in proximity, with the AMB domain oriented parallel to the N-terminal stalk region (Figure S8f). Fits with single models are a crude approximation because in solution, Khc likely is in a dynamic equilibrium of a multitude of autoinhibited and more open conformations. Nevertheless, such single models reflect a possible predominant or average conformation of the protein. These data therefore suggest that binding of Tm1 to Khc results in a shift of the conformational equilibrium towards different, more compact structures of the Khc stalk domain. The autoinhibited conformation of Khc is likely stabilized by interactions within the Tm1-bound stalk in addition to the motor-tail interaction. This is in agreement with our findings that the Khc AMB domain in complex with Tm1 can inhibit Khc independently of the IAK motif.

### Tm1 links inactive Khc to dynein RNPs for minus-end-directed *osk* transport

Khc and Tm1 are loaded onto *osk* RNPs in nurse cells prior to dynein-driven transport into the oocyte^42^. We reasoned that during this process, enhanced inhibition of Khc by Tm1 would favor minus-end-directed movement by preventing a tug-of-war between dynein and Khc that could result in counterproductive movements of RNPs away from the oocyte. To assess if this is the case, we set out to reconstitute *osk* RNPs containing Khc, Tm1 and dynein *in vitro*.

Since dynein and kinesin require different conditions for optimal motility, we used a modified assay buffer that allows movement of both motor classes^58^. To exclude the possibility that Tm1 promotes minus-end-directed movement of *osk* RNPs by stimulating dynein activity directly, we first assessed its effects on the motile properties of the minimal machinery for dynein-mediated RNA transport (dynein, dynactin, BicD, Egalitarian (DDBE)) assembled with an *osk* 3’UTR fragment that is sufficient to initiate dynein movement *in vivo* (*osk* 3’UTR 2+3)^48^. While Tm1 could be observed translocating with dynein and the *osk* RNA (Figure 7a), its presence did not alter dynein velocity or run length on microtubules (Figure 7b, c and Table S4). Furthermore, Tm1 had no effect on the percentage of microtubule-bound dyneins that undergo directed motility or the frequency of dynein binding to microtubules or initiating transport along them (Figure S9a-c and Table S4). Thus, Tm1 does not directly affect dynein motility. Treatment of these reconstituted RNPs with RNase A after assembly dramatically reduced the incidence of Tm1 association with DDBE-*osk* 3’UTR 2+3 (Figure 7d, e), indicating that Tm1 is linked to the dynein machinery via *osk* RNA rather than through protein-protein interactions. This observation is consistent with the finding that Tm1 does not directly influence DDBE activity.

**Figure 7:**
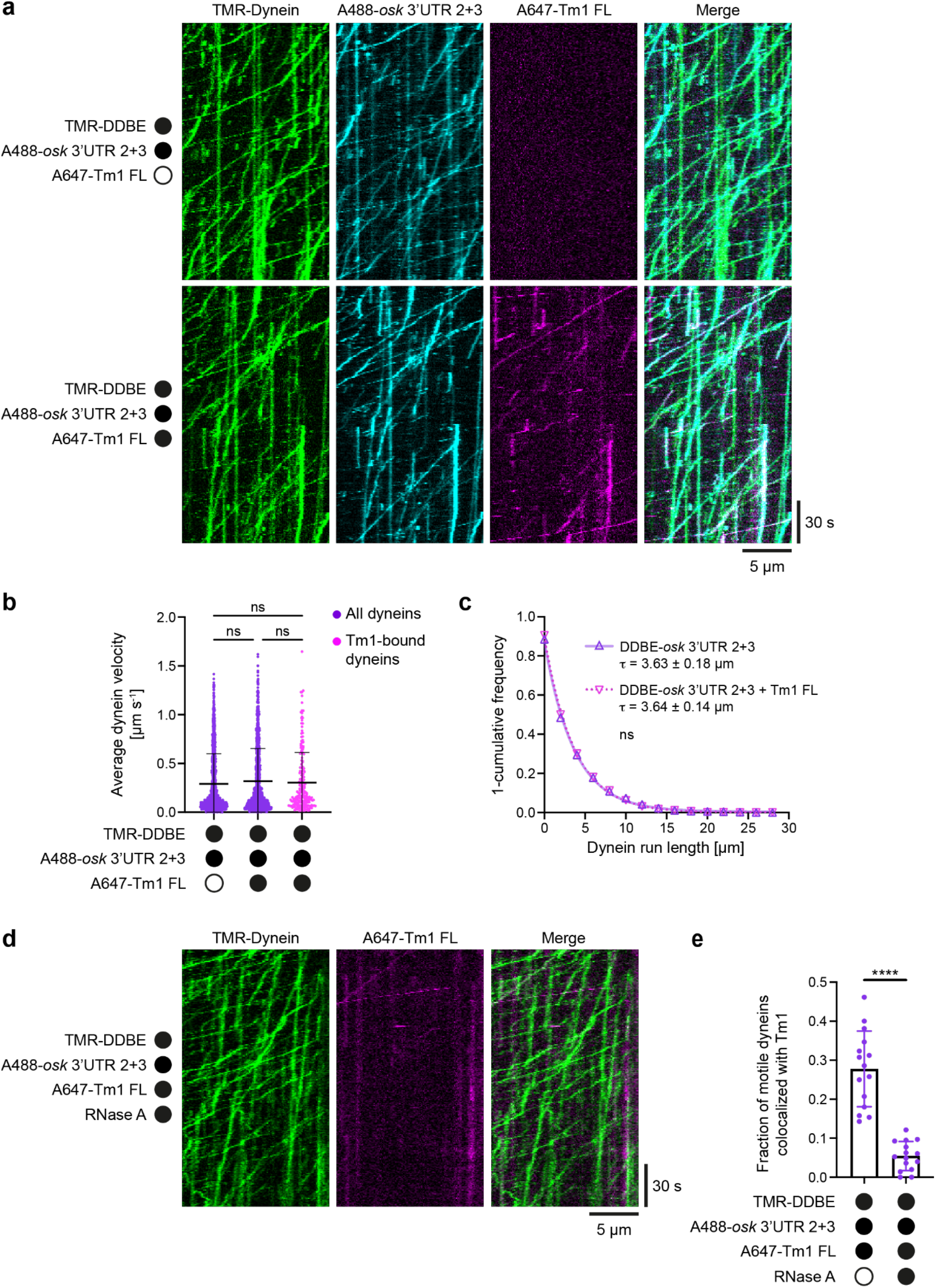
Tm1 does not affect dynein activity in *osk* RNPs. **a** Kymographs showing translocation of dynein and *osk* 3’UTR 2+3 in the presence and absence of Tm1 as part of minimal minus-end-directed RNPs containing dynein, dynactin, BicD, and Egalitarian (DDBE). Co-translocation of Tm1 is observed when present in the assay. **b, c** Average velocities (b) and run lengths (c) of DDBE-*osk* 3’UTR 2+3 complexes assembled with and without Tm1. 1-cumulative frequency distributions of measured run lengths were fit as one-phase decays with the decay constant τ ± 95 % confidence interval shown. Hollow triangles represent empirical bin values used for fitting. **d** Kymographs showing reduced association of Tm1 with motile DDBE-*osk* 3’UTR 2+3 RNPs after treatment with RNase A. **e** Fraction of moving DDBE-*osk* 3’UTR 2+3 RNPs that colocalized with Tm1 signal with and without RNase A treatment. Dots represent values for individual complexes (b) or average values of all motility on individual microtubules (e). Microtubule plus and minus ends are oriented toward the right and left of each kymograph, respectively (a and d). For all plots, mean ± SD is shown. Statistical significance was determined by a Kruskal-Wallis ANOVA test (b), Mann-Whitney test (c), or Welch’s t-test (e). ns: not significant. ****: p<0.0001.

We next assessed the behavior of Khc in the presence of DDBE-*osk* 3’UTR 2+3 RNPs but without Tm1. In these conditions, dynein and associated *osk* RNA exhibited long movements toward the minus ends of microtubules (Figure 8a), as shown by polarity marking of the tracks (Figure S10). Of the relatively few Khc runs that were observed, only ∼25 % were directed towards the microtubule plus end and those were seldomly associated with *osk* RNA (Figure 8a-c and Table S5). The remainder moved towards the minus end in association with dynein and the RNA (Figure 8a-c), indicating some capacity of Khc to associate with DDBE-*osk* RNA complexes. Khc in these complexes appeared to retain some activity, demonstrated by occasional plus-end-directed movements after dissociating from DDBE-*osk* RNPs translocating in the minus end direction (9/127 complexes, 7.1 %; Figure S11). Inclusion of Tm1 in the assay caused a 12-fold increase in the incidence of minus-end-directed Khc transport events (Figure 8a, b and Table S5), which overlapped with dynein and *osk* RNA signals (Figure 8c). These data reveal that Tm1 promotes formation of dual-motor *osk* RNPs in which dynein activity is dominant over that of kinesin, resulting in an overwhelming minus-end-directed bias in Khc movement. Treatment of these *osk* RNPs with RNase A strongly suppressed the Tm1-induced association of Khc with minus-end-directed DDBE- *osk* 3’UTR 2+3 RNA complexes (Figure S12), supporting the notion that Tm1 achieves this effect through interactions with *osk* RNA.

**Figure 8:**
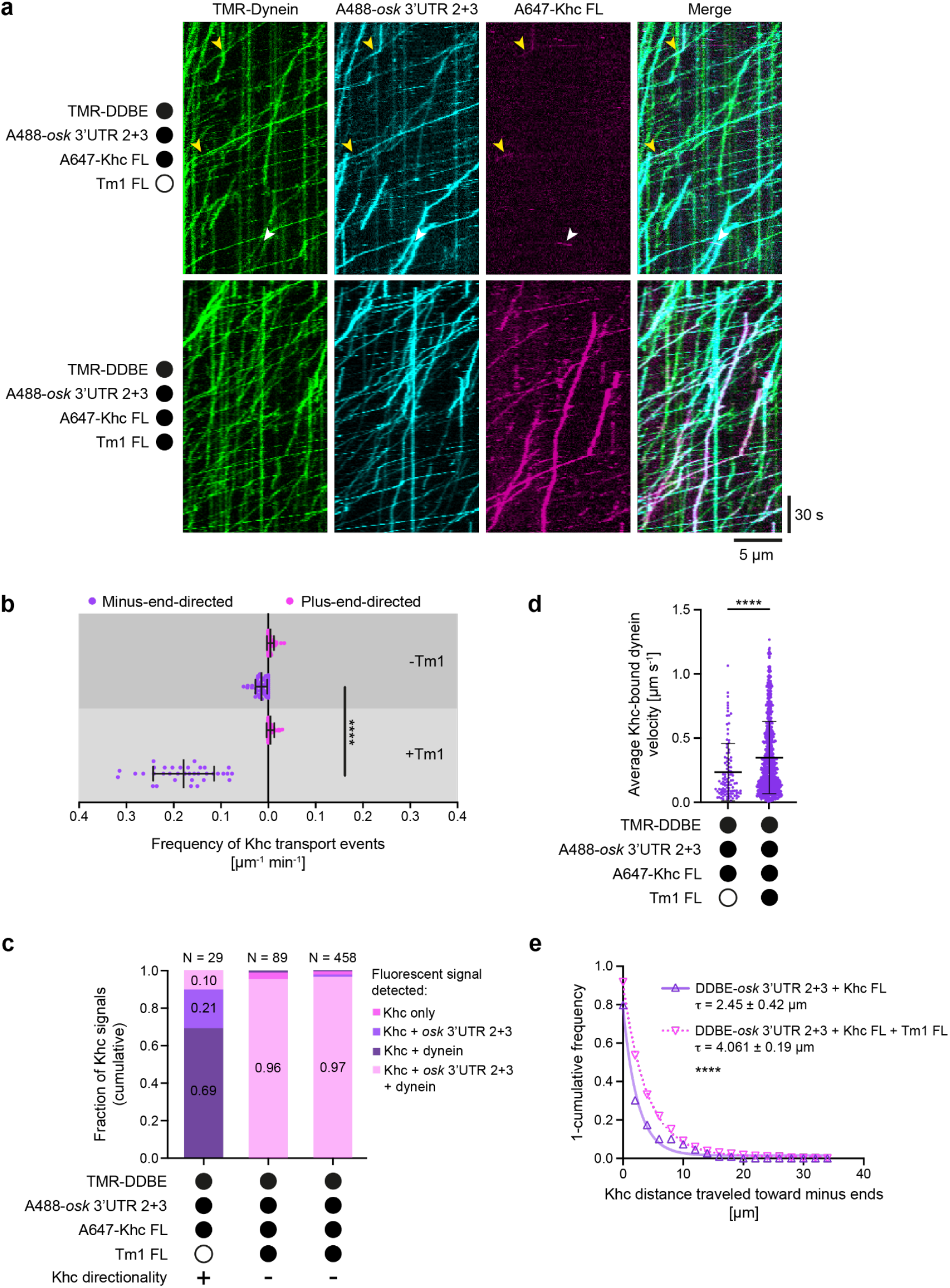
Tm1 promotes association of inhibited Khc with minus-end-directed DDBE-*osk* RNPs. **a** Kymographs showing motile behavior of DDBE (via TMR-dynein), AlexaFluor647-Khc, and AlexaFluor488 *osk* 3’UTR 2+3 in the presence and absence of unlabeled Tm1. Plus- and minus-end-directed Khc movements in the absence of Tm1 are indicated by white and yellow arrowheads, respectively. Microtubule plus and minus ends are oriented toward the right and left of each kymograph, respectively. **b** Frequency of processive Khc movements towards microtubule minus ends (towards left of x-axis, purple) and plus ends (towards right of x-axis, magenta) in the presence and absence of Tm1. **c** Cumulative fraction of plus- and minus-end-directed Khc signals that co-migrated with all combinations of other labeled assay components in the presence and absence of Tm1. **d** Average velocities of minus-end-directed DDBE-*osk* 3’UTR 2+3 complexes with co-migrating Khc signal in the presence and absence of Tm1. **e** Distance traveled by Khc towards the minus end in the presence and absence of Tm1. 1-cumulative frequency distributions of measured run lengths were fit as one-phase decays with the decay constant τ ± 95 % confidence interval shown. Hollow triangles represent empirical bin values used for fitting. Dots represent average values for all motility on individual microtubules (b) or values for individual complexes (d). For all plots mean ± SD is shown. Statistical significance was determined by Welch’s t-test (b) or Mann-Whitney test (d, e). ****: p<0.0001.

To determine if Tm1 additionally suppresses the activity of Khc within minus-end-directed DDBE-*osk* RNPs, we compared dynein velocities and the distances traveled by Khc toward microtubule minus ends in the presence and absence of Tm1. Consistent with an inhibitory effect on Khc activity, Khc-bound DDBE-*osk* RNA complexes moved ∼50 % more rapidly towards the minus end in the presence of Tm1 (Figure 8d and Table S5). Moreover, Tm1 allowed Khc to associate with minus-end-directed DDBE-*osk* RNA complexes for distances on microtubules that were ∼65 % greater than in its absence (Figure 8e and Table S5), and significantly reduced the incidence of Khc directional switches (6/848 complexes, 0.7 %; p<0.0001 compared to in the absence of Tm1 (Fisher’s exact test)). These observations suggest that Tm1 limits engagements of Khc with microtubules that counter minus-end-directed motion. Collectively, these results support a model in which Tm1 recruits Khc to the DDBE-*osk* complex in an inactive state thus avoiding a tug-of-war between the opposite polarity motors.

## Discussion

Tm1-I/C, an atypical RNA-binding isoform of tropomyosin 1, was recently implicated as an RNA adaptor for kinesin-1 transport^41,42^. By interacting directly with Khc’s cargo-binding domain, Tm1 was shown to enhance RNA binding by Khc^43^. Our current study has revealed a previously unknown role of Tm1 in *osk* mRNP transport. We show that Tm1 negatively regulates Khc activity, which we propose occurs by inducing a conformational change in the Khc stalk domain that stabilizes the inhibitory Khc motor-tail interaction (Figure 9a). We demonstrate that Tm1 also links Khc in an inactive state to the dynein transport machinery. This work involves the first reconstitution of a native cargo-adaptor-motor complex that recapitulates regulation of both dynein and kinesin motors. *In vivo*, the mechanism we uncover would allow dynein-mediated transport of *osk* RNPs from the nurse cells to the oocyte to proceed without a tug-of-war with Khc whilst positioning the plus-end-directed motor on the RNPs for when it needs to drive their posteriorward movement within the mid-oogenesis oocyte (Figure 9b).

**Figure 9:**
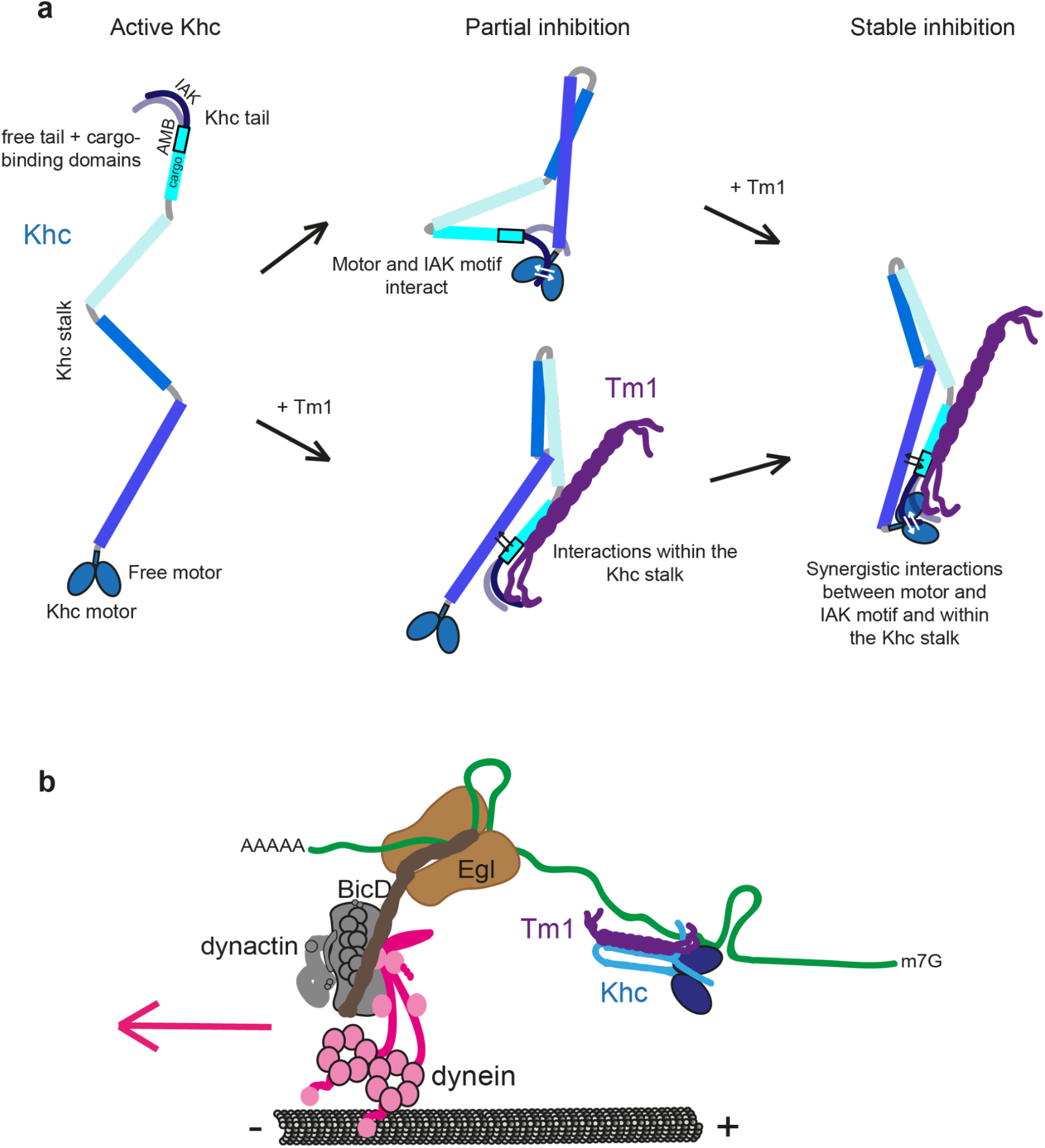
**a** Model of Tm1-mediated stabilization of the auto-inhibited conformation of Khc. Whereas both the motor domain-IAK peptide interaction and interactions within the Tm1-bound, rearranged Khc stalk region, including the AMB domain, can inhibit Khc partially, synergy between these weak interactions stabilizes the inhibiting head-tail interaction. **b** Model of Tm1’s role in dynein-mediated transport of *osk*. By linking Khc to the dynein-transported *osk* RNA and keeping Khc inhibited, Tm1 allows the first, dynein-mediated transport step from the nurse cells to the oocyte to proceed without a tug-of-war between the motors. These events also position Khc on the RNP for the second, Khc- mediated transport step.

### Kinesin-1 autoinhibition mechanisms

Despite decades of research, kinesin-1 autoinhibition is not fully understood. This is due in part to a paucity of structural information for full length kinesin-1. Whereas crystal structures of kinesin motor domains in different states are available, the conformations of the coiled-coil stalk in both the active and autoinhibited state of the motor remain elusive. While an extended conformation of the coiled-coil stalk was proposed to exist in active kinesin-1^59^, this notion was challenged by subsequent studies that indicated an at least partially folded structure of the tail within the active complex^21,60^. A recent study involving structure prediction and single-particle negative stain electron microscopy questions if folding occurs at the flexible hinge region 2 and suggests a different conformation for the Khc stalk^61^.

It has been proposed that the interaction of the IAK motif within the tail with the motor domain plays a key role in kinesin-1 autoinhibition. Consistent with this notion, we observed hyperactivity of *Drosophila* Khc motility when the IAK motif was deleted. However, it was recently shown that Klc inhibits Khc activity independently of the IAK-motor interaction^22^. We also find that the IAK motif is not needed for inhibition of Khc by Tm1. Instead, we find that the Khc AMB domain, a region outside the IAK motif previously shown to be essential for *osk* mRNA localization^62^, contributes to inhibition in Khc-Tm1 complexes. Our work therefore identifies a novel role for the AMB domain.

Using NMR spectroscopy, we show that the IAK-motor interaction does not interfere with binding of regulating factors such as Tm1. Our SAXS data-driven structure modeling provides insights into possible conformations of a full length kinesin-1. The Khc stalk adopts different conformations when in complex with Tm1 (Figure 6c, d). This could explain why the IAK-motor domain interaction, despite being necessary for autoinhibition, is insufficient to fully inhibit Khc. It is conceivable that additional interactions in the rearranged, Tm1-bound stalk regions are required to maintain the autoinhibited, folded conformation. This is in line with our own and previous observations that the IAK-motor domain interaction is rather weak^17^ and the report in a recent preprint that interactions within the coiled-coil stalk and between stalk and motor contribute to kinesin-1 autoinhibition^63^. Presumably, binding of Tm1 drives structural rearrangements of the stalk that compact the molecule and stabilize the autoinhibited conformation to robustly shut down Khc motility. We propose a model in which several weak interactions – the motor-IAK interaction and interactions within the Tm1-bound coiled-coil stalk, including the AMB domain – act synergistically to achieve this stable inhibited conformation of Khc *in vivo* (Figure 9a).

With its functions in cargo-binding and motor regulation, we speculate that Tm1 acts as an alternative Klc in the *Drosophila* germline to regulate processes which were previously thought not to require a Klc^40^. Previously, PAT1 was shown to act as a positive regulator of Khc and have redundant functions with Klc in the *Drosophila* germline ^39^. As correct RNA localization during oogenesis is critical for development, positive regulators such as PAT1 and negative regulators such as Tm1 may have replaced Klc in order to provide nuanced regulation of Khc activity. In the future, it will be interesting to determine whether Klc is also dispensable in other tissues that use Tm1 for RNA localization, including the follicle cells of the egg chamber^64^ and the larval neuromuscular junction^65^.

### Co-ordination of kinesin-1 and dynein motors on *osk* RNPs

Although many cargo types have been shown to be transported by the action of dynein and kinesin-1 motors, the mechanisms of motor regulation in the context of dual motor transport have been elusive. We have identified one of very few known examples of a factor that links dynein and kinesin-mediated transport. Furthermore, to our knowledge, Tm1 is the first example of a protein that modulates dynein and kinesin behavior in a reconstituted dual-motor assembly on a native cargo.

Several studies have reported that active dynein and kinesin motors engage in a tug-of-war when artificially coupled^66,67,68,69,70,71^. However, the relevance of such mechanical antagonism in localizing physiological cargoes for these motors is unresolved^6^. Although we also observe motor opposition in reconstituted RNPs containing DDBE and Khc, our observation that Tm1 supports efficient dynein-mediated RNA localization through robust inhibition of Khc highlights the importance of regulatory factors in addition to mechanical coupling in cargo-adaptor-motor complexes.

Despite the potential for antagonistic engagement of dynein and kinesin on shared cargoes, the motors can exhibit a puzzling co-dependence, in which strongly impairing the activity of one motor type can inhibit the activity of the other^6^. Nonetheless, enhanced inhibition of Khc by Tm1 in our reconstituted RNPs demonstrates the importance of negative kinesin regulation in enabling robust dynein activity. Supporting the *in vivo* relevance of this regulatory logic, kinesin-1 activating IAK mutations were recently shown to impair dynein-mediated transport processes in *Aspergillus nidulans*^72^. Collectively, these observations point to complex interplay between opposite polarity microtubule motors during bidirectional transport. Further reconstitutions of dual-motor systems on native cargoes should reveal generalities of dynein-kinesin crosstalk, as well as any cargo-specific mechanisms.

### Potential kinesin-1 activation mechanisms in the female germline

Our study provides mechanistic insight into two critical aspects of *osk* mRNA transport – the assembly of the dual motor complex and how Khc activity is suppressed during dynein-mediated delivery of the transcript from the nurse cells to the oocyte. However, it is not known how Tm1-mediated inhibition of Khc is alleviated at later stages of oogenesis in the oocyte. One candidate to fulfill this role is Ensconsin. This protein is required for posterior *osk* localization^73^, as well as organelle transport in *Drosophila* neurons^74^ and is homologous to MAP7, which was recently shown to be required for full activity of the human kinesin-1 complex *in vitro*^22,75^. Other factors that may relieve Khc inhibition in the oocyte include the exon junction complex (EJC), which, together with the SOLE RNA structure, is essential for *osk* transport to the posterior of the oocyte^46,76^. As Tm1 remains bound to the *osk* mRNP throughout transport and is required for Khc-mediated translocation of the mRNA to the posterior of the oocyte^42,43^, it is likely that the activating factor(s) induces a conformational change in the Khc-Tm1 complex rather than dissociation of Tm1. Investigating the molecular mechanisms that underpin these dynamics is likely to inform our understanding of how kinesin-1 activity is orchestrated in other transport systems.

## Materials and Methods

### Molecular Cloning

Khc FL for *in vitro* studies was cloned with a HisSUMO-SNAP-3C-tag into MCS1 between BamHI and HindIII sites of the pFastBacDual vector by InFusion cloning (Takara). The Khc mutant constructs used for *in vitro* studies were generated by site-directed mutagenesis of the Khc FL plasmid using primers that either inserted a stop codon after aa 910 (Khc910) or aa 940 (Khc940) or deleted aa 911-938 (KhcΔAMB) followed by DpnI digest of the template plasmid and transformation of the product in *E. coli* XL1-blue cells. For co-expression of Khc FL and Tm1, the Tm1 coding sequence was inserted into MCS2 of pFastBacDual-HisSUMO- SNAP-3C-Khc between SmaI and KpnI sites by InFusion cloning (Takara). Tm1 FL for motility assays was cloned with a HisSUMO-SNAP-tag between SacI and HindIII sites of the pET11 vector by InFusion cloning (Takara). The SNAP-Tm1 1-335 truncation was made by site-directed mutagenesis of the SNAP-Tm1 FL plasmid using primers that inserted a stop codon after aa 335 as described above.

### Protein expression and purification

His6-SUMO-SNAP-3C-tagged Khc FL, Khc910 or ΔIAK and ΔAMB truncations and co-expressed Khc-Tm1 complexes were expressed using the Bac-to-Bac Baculovirus expression system (Gibco). 1 L of *Sf21* insect cells at 1x10^6^ cells mL^-1^ shaking culture were infected with 10 mL of P1 virus stock. After expression for 3 d at 27.5 °C, cells were harvested by centrifugation at 1000 g for 20 min at 4°C. The pellets were lysed in a dounce tissue grinder in lysis buffer (20 mM Tris-HCl at pH 7.5, 500 mM NaCl, 1 mM MgCl_2_, 0.1 mM ATP, 2 mM DTT, 5 % glycerol, 0.2 % Tween-20). The lysates were cleared by centrifugation and the soluble protein fraction was affinity-purified on a HisTrap Excel column (GE Healthcare). After elution in a 0–300 mM imidazole gradient, the HisSUMO-fusion tag was cleaved either by Senp2 (SNAP-tagged proteins for fluorescent labeling) or 3C protease (unlabeled proteins) digest during dialysis in GF150 buffer (25 mM HEPES/KOH pH 7.3, 150 mM KCl, 1 mM MgCl_2_, 0.1 mM ATP, and 2 mM DTT) for 16 h before further purification by anion exchange chromatography on a HiTrap Q HP column (GE Healthcare), followed by size exclusion chromatography (SEC) on a Superdex 200 10/300 Increase column (GE Healthcare).

His6-SUMO-tagged and His6-SUMO-SNAP-tagged Tm1 protein constructs and GST-Khc 1- 365 were expressed in BL21-CodonPlus(DE3)-RIL cells (Stratagene) by isopropyl β-D-1- thiogalactopyranoside (IPTG) induction for 16 h at 18 °C. Cells were grown in Luria-Bertani (LB) medium (0.5 % [w/v] yeast extract [MP], 1 % [w/v] tryptone [MP], 0.5 % [w/v] NaCl at pH 7.2) supplemented with antibiotics (100 μg mL^-1^ ampicillin or 10 μg mL^-1^ kanamycin and 34 μg mL^-1^ chloramphenicol). After harvesting, the pellets were lysed by a microfluidizer processor (Microfluidics) in 20 mM Tris-HCl (pH 7.5), 500 mM NaCl, 5 % glycerol, 0.01 % NP- 40, and 40 mM imidazole buffer supplemented with protease inhibitor cocktail (Roche) and 2 mM DTT. The lysates were subsequently cleared by centrifugation. HisSUMO-tagged proteins were affinity-purified by Ni-IMAC on a HisTrap column (Cytiva) and eluted with an imidazole gradient (40–300 mM). After cleavage of the His6-SUMO-tag with Senp2 protease, cleaved tags and protease were removed by a second, subtractive Ni-IMAC on a HisTrap column (GE Healthcare). The proteins were further purified on either a 10/300 Superose 6 Increase GL or a HiLoad 16/600 Superdex 200 SEC column in either GF150 buffer for motility assays or NMR buffer (25 mM NaPO_4_ pH 6.5, 150 mM NaCl, and 0.5 mM TCEP) for NMR experiments.

GST-tagged Khc 1-365 was affinity-purified on a GSTrap column (Cytiva) and glutathione gradient (1-20 mM). The protein was further purified on a HiLoad 16/600 Superdex 200 SEC column in GF150 buffer.

SNAP-tagged Khc and Tm1 proteins were fluorescently labeled with SNAP-Cell TMR Star, SNAP-Surface Alexa Fluor 488 or SNAP-Surface Alexa Fluor 647 at 15 °C for 90 min in GF150 buffer with a 1.5x excess of dye. Free dye was removed by a subsequent desalting step on a PD-10 desalting column (Cytiva). Labeling efficiencies were determined by measuring protein concentration at 280 nm and dye concentration at the dye’s excitation maximum. For monomeric protein, labeling efficiency was calculated as c_dye_/c_protein_. For dimeric proteins, labeling efficiency was calculated based on monomer labeling efficiencies as (L_1_ x U_2_) + (U_1_ x L_2_) + (L_1_ x L_2_) with L1 = fraction of monomer 1 labeled, L2 = fraction of monomer 2 labeled, U1 = fraction of monomer 1 unlabeled, U2 = fraction of monomer 2 unlabeled.

Recombinant human dynein and *Drosophila* Egl-BicD complexes were expressed in *Sf9* insect cells with the dynein complex fluorescently labeled via a SNAPf tag fused to the N-terminus of Dynein heavy chain, as described previously^15^. Native porcine dynactin was purified from brain tissue as described previously^15^.

### RNA *in vitro* transcription

The *osk* RNA constructs used for the motility assays were synthesized *in vitro* using the MEGAscript T7 Transcription Kit (Ambion). The RNA was transcribed from a PCR product amplified with a T7 promote-containing forward primer and containing an *osk* mRNA FL, *osk* 3’UTR or a ∼529-nt region of the *osk* 3’UTR previously defined as region 2+3 (*osk* 3’UTR 2+3) that is sufficient to promote the localization of reporter transcripts to the developing oocyte^48^. The template for *osk* min was generated from three overlapping primers by PCR and subsequently amplified again with a T7 promoter containing forward primer. RNA was transcribed from the final PCR product. For fluorescent labeling of the *osk* 3’UTR and *osk* min transcripts, Cy3- or Cy5-UTP was included in the transcription reaction with a 4-fold excess of unlabeled UTP, yielding transcripts with several labels per RNA molecule. For *osk* 3’UTR 2+3, transcripts were fluorescently labeled with AlexaFluor488-UTP with a 5-fold excess of unlabeled UTP. Transcripts were subsequently purified by phenol/chloroform extraction and passed through a G50 mini Quick Spin RNA desalting column (Sigma-Aldrich) to remove unincorporated nucleotides before sodium acetate/ethanol precipitation. The precipitated RNA was resuspended in nuclease-free water and stored at -20 °C or -70 °C.

### Polymerization and stabilization of microtubules

Microtubules were polymerized from porcine tubulin (Cytoskeleton Inc., Denver, CO) and labeled with fluorophores and biotin by stochastic incorporation of labeled dimers into the microtubule lattice. Mixes of 2.5 µM unlabelled tubulin, 0.17 µM Hilyte488- or AMCA-tubulin, and 0.3 µM biotin-tubulin were incubated in BRB80 (80 mM PIPES pH 6.85, 2 mM MgCl_2_, 0.5 mM EGTA, 1 mM DTT) with 0.67 mM GMPCPP (Jena Bioscience, Jena, Germany) overnight at 37 °C. Polymerized microtubules were pelleted in a room temperature table top centrifuge at 18,400 x g for 8.5 min, and washed once with pre-warmed (37 °C) BRB80. After pelleting once more, the microtubules were gently resuspended in 300 µL pre-warmed (37 °C) BRB80 containing 20-40 µM paclitaxel (taxol; Sigma-Aldrich).

Polarity marked microtubules with HiLyte 488 tubulin enriched at minus ends were made according to the principle described previously^77^. Bright microtubule seeds were polymerized at 37 °C for 10 minutes in a mix containing 2.5 µM unlabelled porcine tubulin (5-fold excess over fluorescent tubulin), 0.5 µM HiLyte 488 tubulin, and 0.3 µM biotin-tubulin in BRB80 with 0.67 mM GMPCPP. After 10 minutes, this mixture was diluted 1:1 with a tubulin polymerization mix containing 12.5 µM unlabelled tubulin and 0.3 µM biotin-tubulin in BRB80 with 0.67 mM GMPCPP for final tubulin concentrations of 7.5 µM unlabelled tubulin (30-fold excess over fluorescent tubulin), 0.25 µM HiLyte 488 tubulin, and 0.3 µM biotin-tubulin. The final tubulin mix was incubated at 37 °C for an additional 2 h to allow growth of relatively dim microtubule plus ends before harvesting and taxol stabilization as described above.

### *In vitro* single-molecule reconstitution assays, TIRF microscopy and image analysis

TIRF-based single-molecule reconstitution assays were performed as described previously^15^. Briefly, taxol-stabilized porcine microtubules were immobilized in imaging flow chambers by streptavidin-based linkages to biotin-PEG passivated cover slips. Khc RNP assay mixes were assembled at 500 nM of each component. Assemblies of the dynein transport machinery with Khc and Tm1 contained 100 nM dynein, 200 nM dynactin, 500 nM Egl-BicD, 1 μM RNA, 500 nM Khc (or Khc storage buffer as a control) and 2 μM Tm1 (or Tm1 storage buffer as a control), and were incubated on ice for 1-2 h prior to imaging. The RNP assay mixes were then diluted to concentrations suitable for imaging of single molecules (6.25 nM for Khc RNP experiments, 5 nM dynein and 25 nM Khc for experiments containing both motors) in BRB12 (Khc experiments) or modified dynein motility buffer^58^ (30 mM HEPES-KOH pH 7.0, 50 mM KCl, 5 mM MgSO_4_, 1 mM EGTA pH 7.5, 1 mM DTT, 0.5 mg mL^-1^ BSA - experiments including DDBE) supplemented with 1 mg mL^-1^ α-casein, 20 μM taxol, 2.5 mM MgATP and an oxygen scavenging system (1.25 μM glucose oxidase, 140 nM catalase, 71 mM 2-mercaptoethanol, 25 mM glucose final concentrations) to reduce photobleaching, and applied to the flow chamber for imaging by TIRF microscopy.

For assays in which RNase A was added, assemblies were diluted into modified dynein motility buffer as described above with either 1 µL of 0.2 µg/mL RNase A (final concentration of 9.5 ng/mL) diluted in motility buffer, or 1 µL of motility buffer alone added in addition. Such low concentrations of RNase A were used to ensure mild digestion of the RNA (which was monitored during acquisition via the fluorescent RNA signal) that would eliminate any RNA-based connections between protein components whilst limiting digestion of the predominantly double-stranded localization signal important for activating DDBE motility^48^. These mixes were incubated at room temperature for 5 min before application to the imaging chambers.

For each chamber, a single multi-color acquisition of 500 frames was made at a frame rate of ∼2 frames s^-1^ and 150 ms exposure (1-2 colors) or 1.2-1.4 frames s^-1^ and 50 ms exposure (3 colors). Images of Khc-Tm1 were acquired using a Leica GSD TIRF microscope system. All images involving the dynein machinery were acquired on a Nikon TIRF system using Micromanager control software^78^ as described previously^15^.

Binding and motility of RNP components in the *in vitro* reconstitution assays were manually analyzed by kymograph in FIJI^79^ as described previously^15^. Statistical analysis and plotting were performed using GraphPad Prism version 9.1.1 for MacOS, (GraphPad Software, San Diego, USA).

As we noticed some variability of Khc activities between different assays, we repeated all appropriate control conditions for each experiment on the same day and used these controls as comparison for our analysis. Results were confirmed on multiple days in duplicates or triplicates of the experiments.

### Microtubule binding assays

For microtubule binding assays, microtubules were polymerized from 5 mg mL^-1^ porcine tubulin (Cytoskeleton Inc., Denver, CO) at 37 °C for 20 min in BRB80 supplemented with 1 mM GTP and then diluted 1:10 in BRB80 + 20 µM taxol. Proteins of interest were incubated at indicated concentrations with 20 µL of this microtubule solution for 30 min at RT in a final volume of 50 µL, before soluble and microtubule-bound fractions were segregated by ultracentrifugation at 80,000 g for 30 min over a 500 µL 30 % sucrose cushion. Soluble fractions were supplemented with 1x SDS-loading dye. Microtubule pellets were washed with BRB80 + taxol before redissolving in 50 µL 1x SDS-loading dye. Soluble and pellet fractions were analyzed by SDS-PAGE.

### GST-pulldown competition assay

To test whether Tm1 competes with the Khc motor-tail interaction, 5 µM GST-Khc 855-975 as bait and an indicated excess of prey protein Khc 1-365 were incubated with increasing concentrations of Tm1 1-335 as competitor for Khc 855-975 binding in a total volume of 100 µL in GF150 buffer (25 mM HEPES/KOH pH 7.3, 150 mM KCl, 1 mM MgCl_2_, 2 mM DTT). After incubation for 40 min on ice, the mixture was added to pre-equilibrated 50 µL Glutathione Sepharose 4B beads (GE Healthcare) and proteins were allowed to bind for 2 h at 4 °C on a nutating shaker. Beads were washed with 4 x 500 µL and 1 x 100 µL GF150 + 0.01 % NP-40. Bound proteins were eluted from the beads by boiling in 1x SDS-loading dye for 5 min at 95 °C. Input, wash and elution samples were analyzed by SDS-PAGE.

### Mass photometry

Mass photometry data were acquired on a ReFeyn OneMP from co-expressed, co-purified Khc-Tm1 complexes at a sample concentration of 20 nM in PBS. Data acquisition was performed using AcquireMP, with data analysis performed in DiscoverMP (Refeyn Ltd, v1.2.3).

### NMR spectroscopy

NMR measurements were performed at 298 K on a Bruker Avance III NMR spectrometer with magnetic field strengths corresponding to proton Larmor frequencies of 600 MHz equipped with a cryogenic triple resonance gradient probe head. NMR sample concentration was 50 µM for titration experiments and 177 µM for backbone assignment experiments. Experiments for backbone assignments were performed on ^13^C,^15^N-labeled samples using conventional triple- resonance experiments (HNCO, HNCA, CBCA(CO)NH, HN(CO)CA and HNCACB)^80^. All spectra were acquired using the apodization weighted sampling scheme^81^ and processed using NMRPipe^82^. Resonance assignments were done with the program Cara.

For titrations, RNA and Tm1 were added to ^15^N-labeled Khc 855-975 at indicated ratios and a ^1^H,^15^N-HSQC was recorded for each titration point. Peak intensity ratios were derived using NMRView^83^ and corrected for dilution. The extent of amide ^1^H-^15^N chemical shift perturbations (CSPs) in free versus bound Khc 855-975 were calculated according to Williamson^84^ to compensate for frequency range differences between ^15^N and ^1^H dimensions.

### SAXS data acquisition and structure modeling

SAXS data on full length Khc in isolation and in complex with full length Tm1 in 25 mM HEPES/KOH pH 7.3, 150 mM KCl, 1 mM MgCl_2_ and 2 mM DTT at concentrations of 0.5, 1.0 and 2.0 mg mL^-1^ were recorded at 20 °C on BM29 beamline, ESRF Grenoble, France. Ten frames with an exposure time of 1 s/frame were recorded in batch mode using an X-ray wavelength of 0.992 Å and a sample distance of 2.81 m to the Pilatus2M detector. Data reduction and buffer subtraction was performed by the automatic processing pipeline^85^. *R_g_* and *I(0)* were determined using the Guinier approximation using Primus/autorg^86^.

The scattering curve was subsequently compared to pools of structures generated by connecting AlphaFold2^55,56^ models of the Khc head domain (residues 1-378) homodimer connected to four stretches of coiled-coil homodimers (residues 403-585, 604-706, 711-838 and 844-936 respectively). The linker regions between the structured parts were modified by random phi/psi rotations in monomer A of the homodimer followed by a short four-step simulated annealing energy minimization to restore proper bond geometries in monomer B, for which all corresponding residues were rotated together with monomer A during the linker randomization. The C-termini of monomer A and monomer B were subsequently randomized independently. A total of 5000 structures were generated in this way. In a second step, the structure of the motor homodimer was replaced by a homodimer (residues 1-344) bound to a peptide from the C-terminus of molecule A (residues 934-953) containing the IAK motif, which was followed by simulated annealing energy minimization to restore proper bond geometry. During the simulated annealing energy minimization steps, the structures of the motor and coiled-coil models or motor/peptide were unaltered by using a harmonic energy term to keep the coordinates close to the starting coordinates in the template structure^87^. The calculations were performed using CNS1.2^88^ within the ARIA1.2 framework^89^. Structure pools for the Khc homodimer in complex with Tm1 were generated in a similar way with an AlphaFold2-modeled α-helix of Tm1 (residues 258-386) bound to the last coiled-coil stretch of Khc (residues 844- 936) in a position derived from the X-ray structure of the Khc-Tm1 complex^43^ and additional randomization of all residues outside the α-helix of Tm1. Each of the 2x5000 models were fitted to the SAXS scattering intensities using CRYSOL^86^.

## Acknowledgements

We thank Lyudmila Dimitrova-Paternoga for reagents, the EMBL Protein Expression and Purification Core Facility, especially Karine Lapouge, the EMBL Advanced Light Microscopy Facility, especially Marko Lampe, and the EMBL Chemical Biology Core Facility, especially David Will, for their support. We thank the Bio-SAXS beamline at European Synchrotron Radiation Facility Grenoble, BM29. We thank Petra Pernot (ESRF), Jérôme Kieffer (ESRF) and Cy Jeffries (EMBL Hamburg) for discussions. S.H. was supported by the EMBL Interdisciplinary Postdoctoral fellowship (EIPOD) Programme under Marie Curie Cofund Actions MSCA-COFUND-FP (664726) and Deutsche Forschungsgemeinschaft (DFG)- Forschergruppe 2333 grant (EP37/4-1) to A.E.. Work in the laboratories of J.H. and A.E. was supported by funding from the DFG via the priority program SPP1935 to J.H. and A.E. (EP37/3- 1 and EP37/3-2) and the EMBL. Work in S.L.B.’s group is supported by the Medical Research Council (MRC), as part of United Kingdom Research and Innovation (also known as UK Research and Innovation) [MRC file reference number MC_U105178790]. M.A.M. is supported by a project grant from the BBSRC (BB/T00696X/1). For the purpose of the MRC open access policy, the authors have applied a CC-BY public copyright license to any Author Accepted Manuscript version arising.

## Author contributions

S.H., M.A.M., A.E., and S.L.B. conceived the study. S.H., M.A.M. and B.S. performed the experiments. S.H., M.A.M., B.S. and J.H. analyzed the data. S.L.B. and A.E. supervised the study. S.H. drafted the manuscript. All authors contributed to the writing of the manuscript.

---

**Figure S1:**
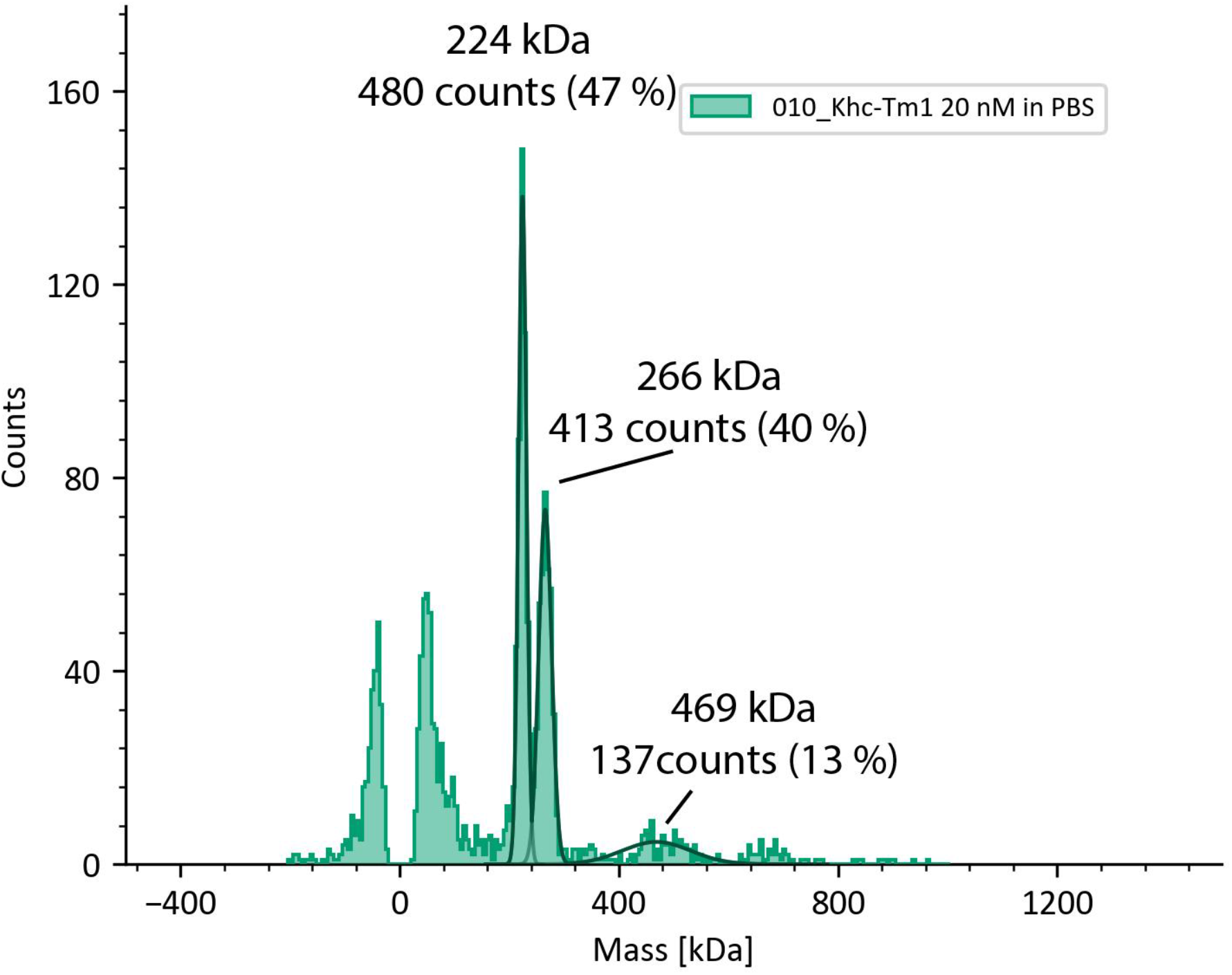
Mass photometry measurements of the Khc-Tm1 complex at 20 nM concentration. The sample comprises a mixture of 47 % particles of 224 kDa, corresponding to the Khc dimer (theoretical MW = 221.2 kDa), 40 % particles of 266 kDa, corresponding to a Khc dimer bound to a Tm1 monomer (theoretical MW = 269.2 kDa), and 13 % of the particles represent larger species. This indicates that in solution, ∼50 % of Khc is bound to Tm1 at steady-state at concentrations below the K_D_, confirming the 2:1 stoichiometry of the Khc-Tm1 complex^43^.

**Figure S2:**
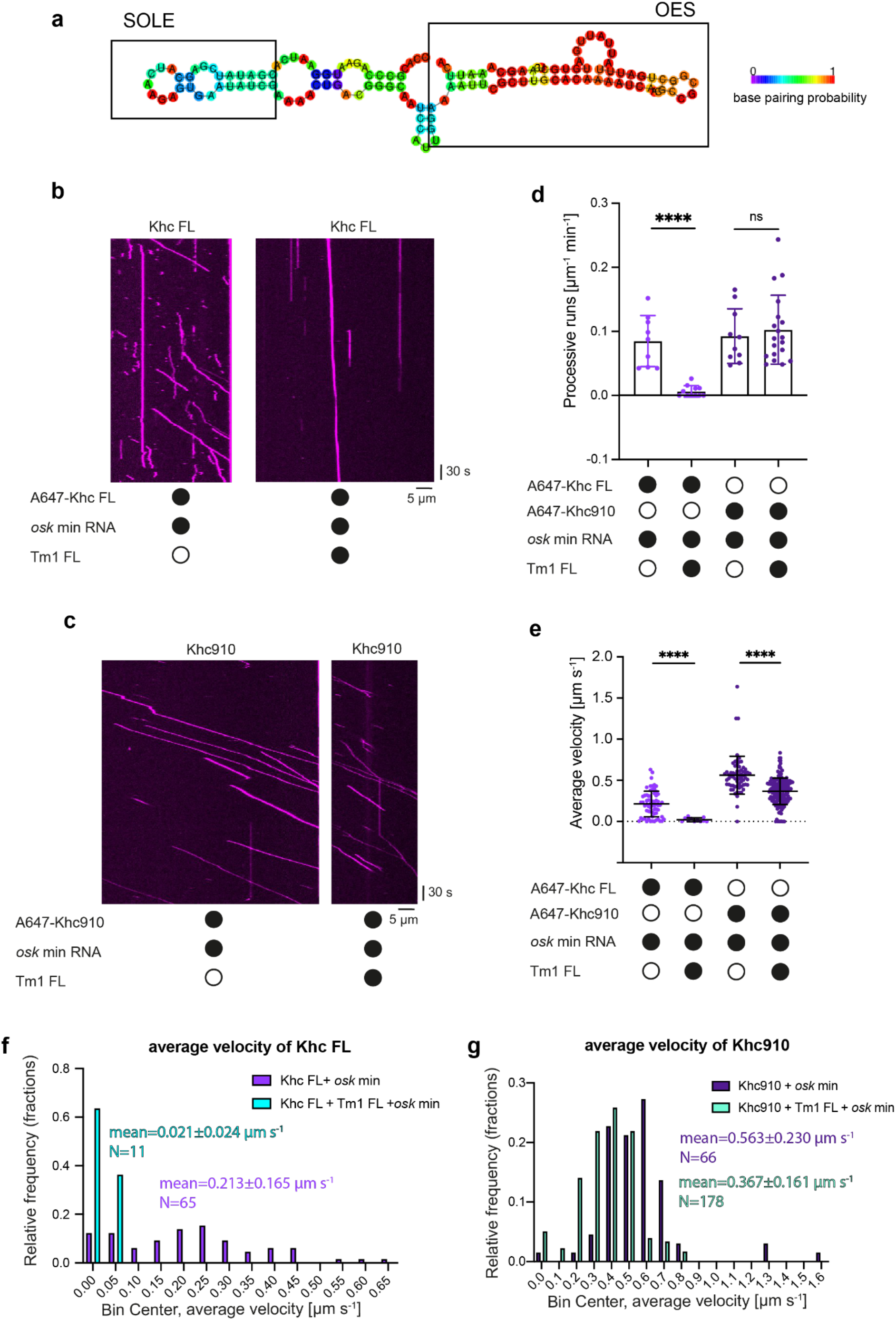
**a** RNAfold ^90^ MFE prediction of *osk* min RNA. Both SOLE and OES are predicted as stem-loop structures, indicating that in the context of *osk* min, the LEs adopt their native folds. **b, c** Example kymographs showing processive movements of AlexaFluor647-labeled SNAP-Khc FL (b) or the tailless Khc910 (c) in absence or presence of Tm1. Microtubule plus and minus ends are oriented toward the right and left of each kymograph, respectively. **d** Processive events of Khc FL and Khc910 in absence or presence of Tm1. In this and other figures, black or white circles indicate proteins that were present or absent from the experiment, respectively. Data points show values for individual microtubules (between 11 and 178 complexes analyzed per condition). Error bars: SD. Statistical significance was evaluated with an unpaired t-test. ****p<0.0001. **e** Average velocities of Khc FL and Khc910 in absence or presence of Tm1. Data points represent individual runs. Statistical significance was evaluated with an unpaired t-test. ****p<0.0001 **f, g** Histograms of the velocity distributions of Khc FL (f) and Khc910 (g) in absence or presence of Tm1.

**Figure S3:**
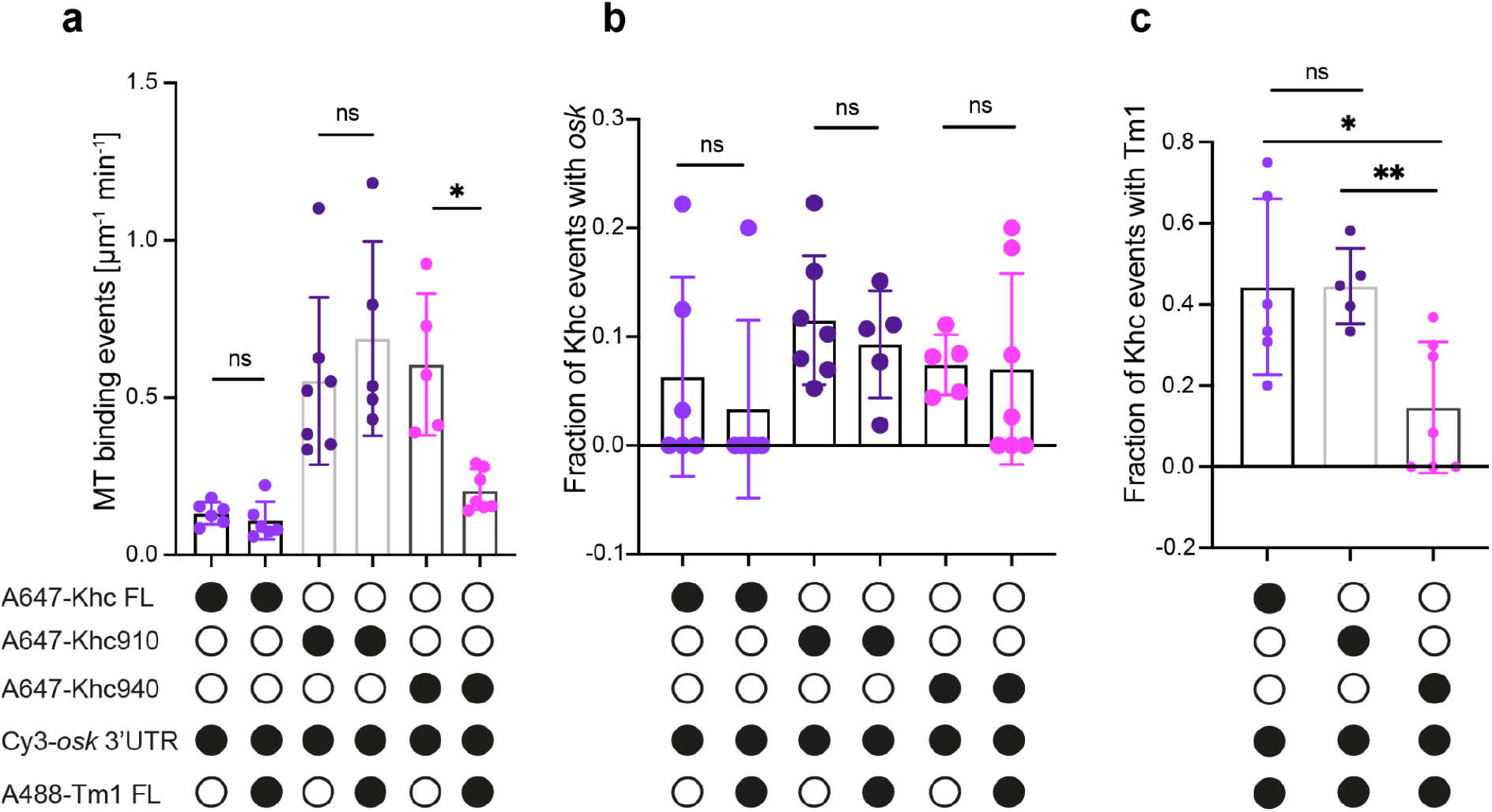
**a** All microtubule (MT)-binding events (processive, static and diffusive) observed for Khc FL, Khc910 and Khc940 ± Tm1 FL. **b** Fraction of observed Khc events colocalizing with *osk* 3’UTR and **c** fraction of observed Khc events colocalizing with Tm1, for Khc FL, Khc910 and Khc940. Data points show values for individual microtubules. Statistical significance was evaluated with an unpaired t-test. *p<0.1 **p<0.01.

**Figure S4:**
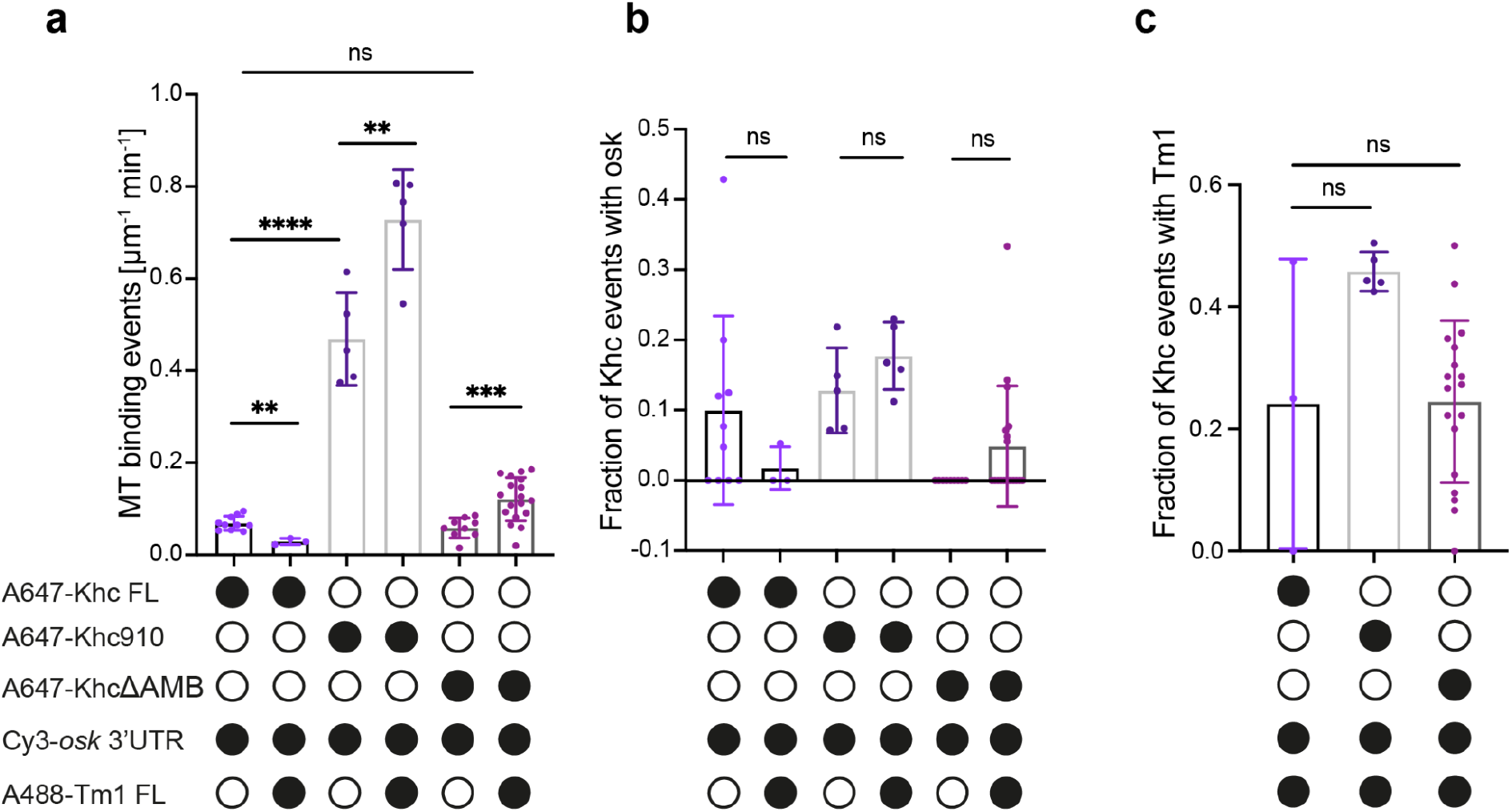
**a** All microtubule (MT)-binding events (processive, static and diffusive) observed for Khc FL, Khc910 and KhcΔAMB ± Tm1 FL. **b** Fraction of observed Khc events colocalizing with *osk* 3’UTR and **c** fraction of observed Khc events colocalizing with Tm1, for Khc FL, Khc910 and KhcΔAMB. Data points show values for individual microtubules. Statistical significance was evaluated with an unpaired t-test. **p<0.01, ***p<0.001, ****p<0.0001.

**Figure S5:**
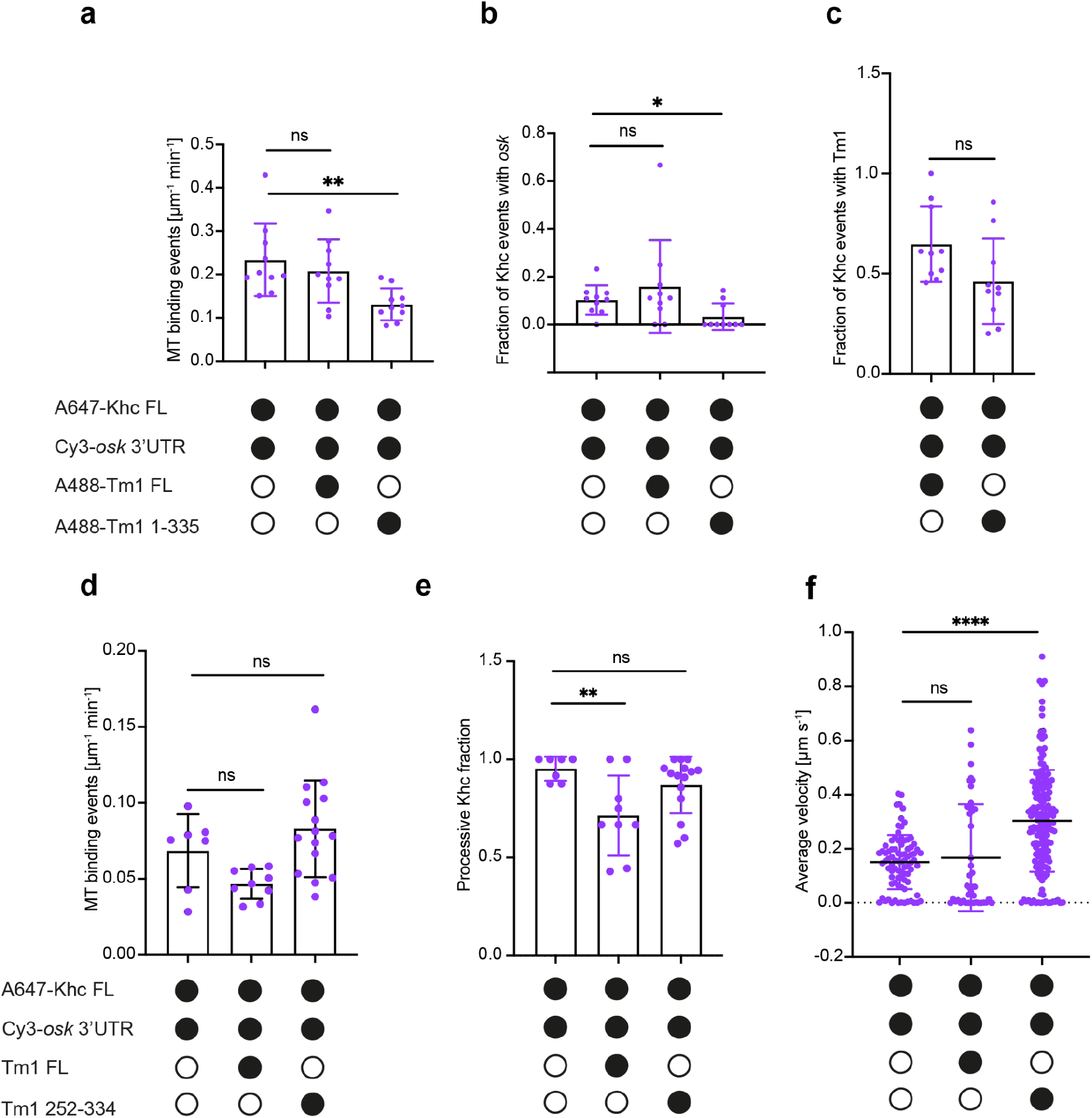
**a** All microtubule (MT)-binding events (processive, static and diffusive) observed for Khc FL ± Tm1 FL or Tm1 1-335. **b** Fraction of observed Khc events colocalizing with *osk* 3’UTR and **c** fraction of observed Khc events colocalizing Tm1 FL or Tm1 1-335. Labeling efficiencies for Tm1 FL and Tm1 1-335 were 57 %, and 58 %, respectively, such that co-localization results are comparable. **d** All microtubule (MT)-binding events (processive, static and diffusive) observed for Khc FL ± Tm1 FL or Tm1 252-334. **e** Fraction of processive events of Khc FL ± Tm1 FL or Tm1 252-334. **f** Average velocities of Khc FL ± Tm1 FL or Tm1 252-334 in the assembly mix. Data points show values for individual microtubules. Statistical significance was evaluated with an unpaired t-test. *p<0.1 **p<0.01 ****p<0.0001.

**Figure S6:**
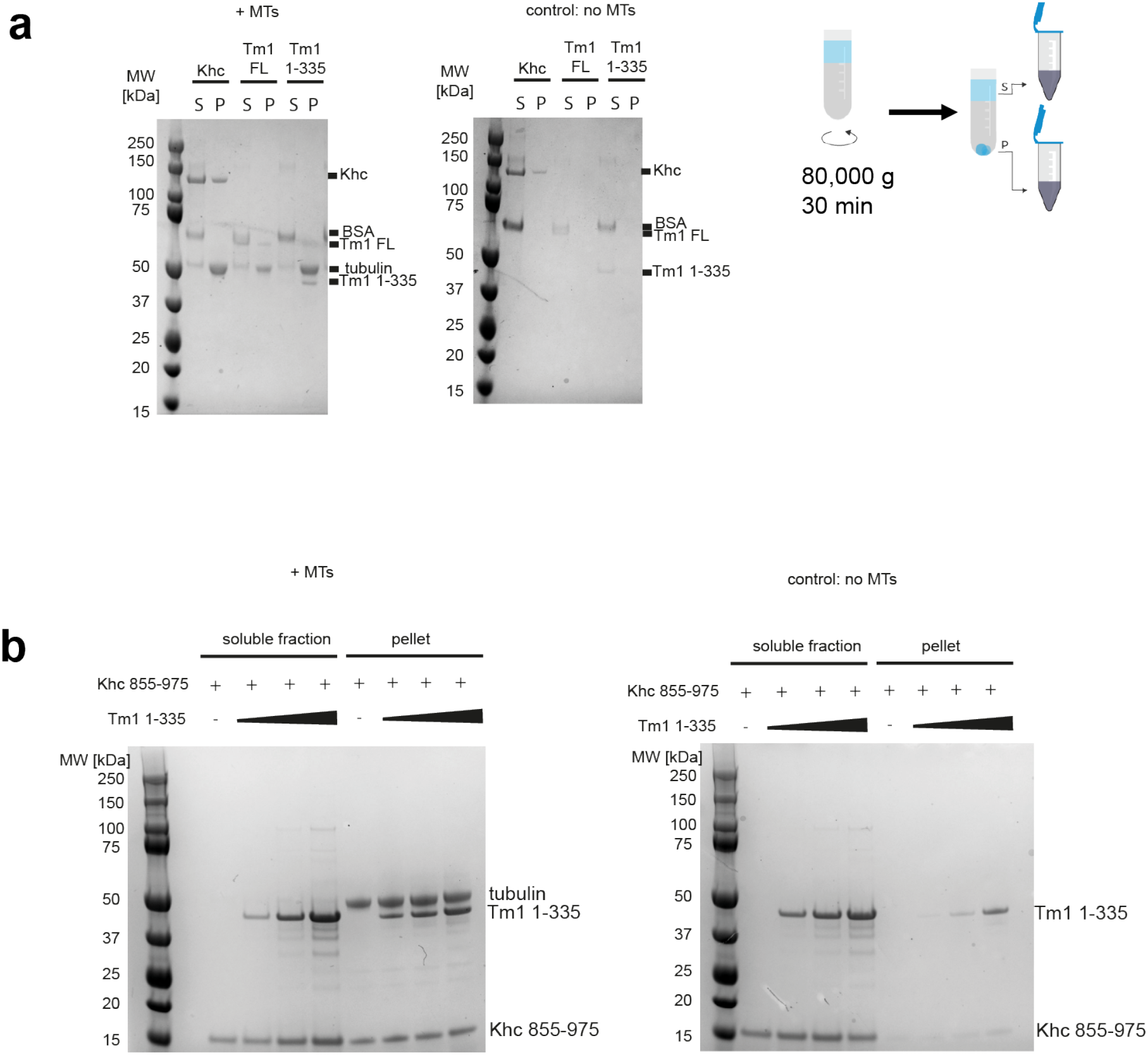
**a** Microtubule binding assay with Tm1 FL and Tm1 1-335. Khc FL served as a positive control. Both Tm1 FL and Tm1 1-335 partition partially into the microtubule pellet fraction. 10 µg of each protein were incubated with polymerized microtubules prior to segregation of soluble and microtubule-bound fractions by ultracentrifugation over a 30 % sucrose cushion. BSA was added to all samples as a negative control. To control for microtubule-independent pelleting of the proteins, controls without microtubules were performed. Soluble and microtubule pellet fractions were analyzed by SDS-PAGE. **b** Microtubule binding assay with Khc 855-975 and Tm1 1-335. Khc 855-975 partially partitions into the microtubule pellet both in absence and presence of increasing concentrations of Tm1 1-335. 25 µM Khc 855-975 were incubated with Tm1 1-335 at ratios 1:0.5, 1:1 and 1:2 and polymerized microtubules before proceeding as described above.

**Figure S7:**
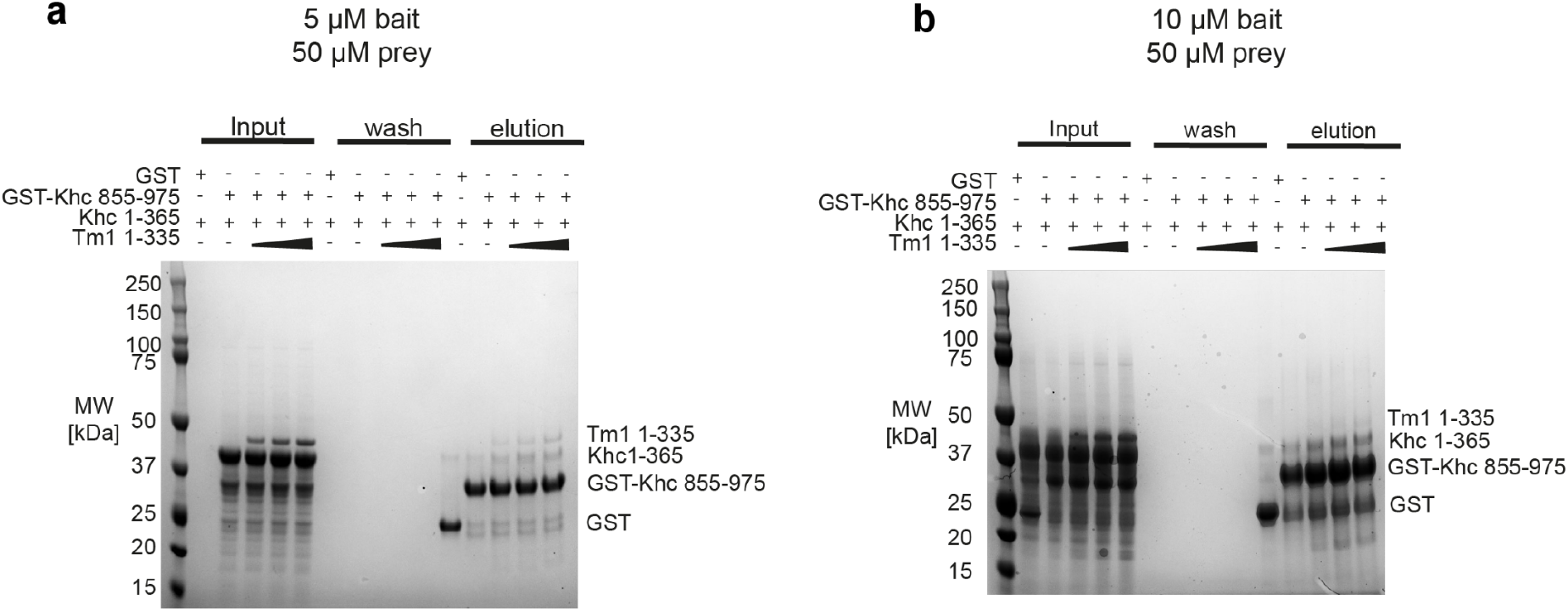
GST-pulldown competition assays to assess the effect of Tm1 on Khc tail - motor interaction. **a** 5 µM or **b** 10 µM GST-Khc 855-975 as bait and 50 µM Khc 1-365 as prey. Tm1 1-335 was added at increasing concentrations (5 to 15 µM) as a competitor for binding to GST-Khc 855-975.

**Figure S8:**
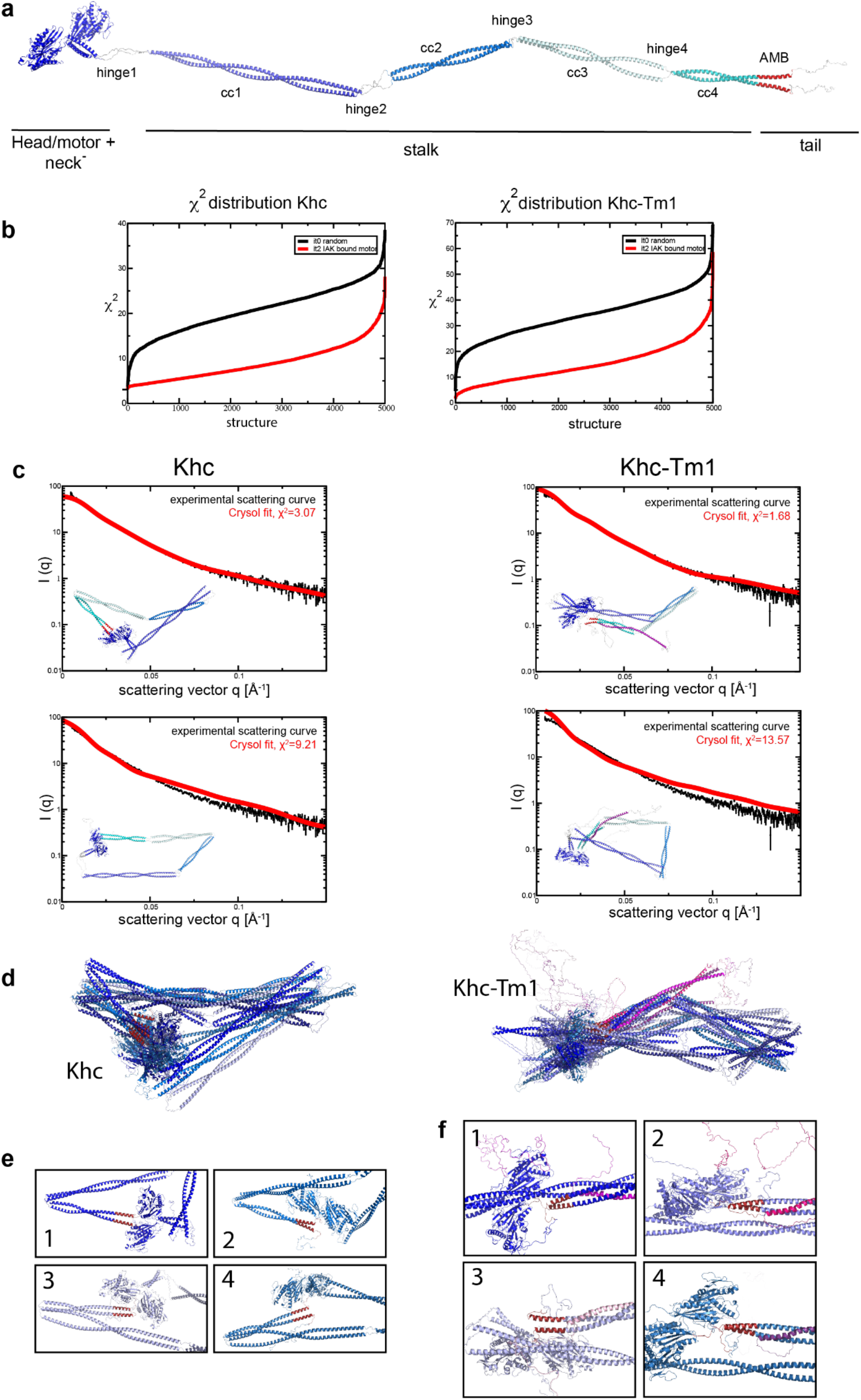
**a** Template model of Khc used for randomization. Coordinates of the structured parts (motor domain, residues 1-378 and the four coiled-coil regions, residues 403-585, 604-706, 711-838 and 844- 936, respectively) were generated by AF2 ^55,56^ and connected with the coordinates of the missing linker residues. **b** χ^2^ distribution of all Crysol fits to the Khc (left) and Khc-Tm1 (right) scattering curves. **c** Best (upper panel) and mediocre (lower panel) fits of Khc (left) and Khc-Tm1 (right) structural models to the experimental scattering curves. **d** Five best fits for Khc (left) and for Khc-Tm1 (right) aligned. **e** Close- ups of the AMB domains in the four best fitting models for Khc. **f** Close-ups of the AMB domains in the four best fitting models for Khc-Tm1. Khc models are shown in shades of blue with the AMB domains marked in red, Tm1 is shown in shades of pink.

**Figure S9:**
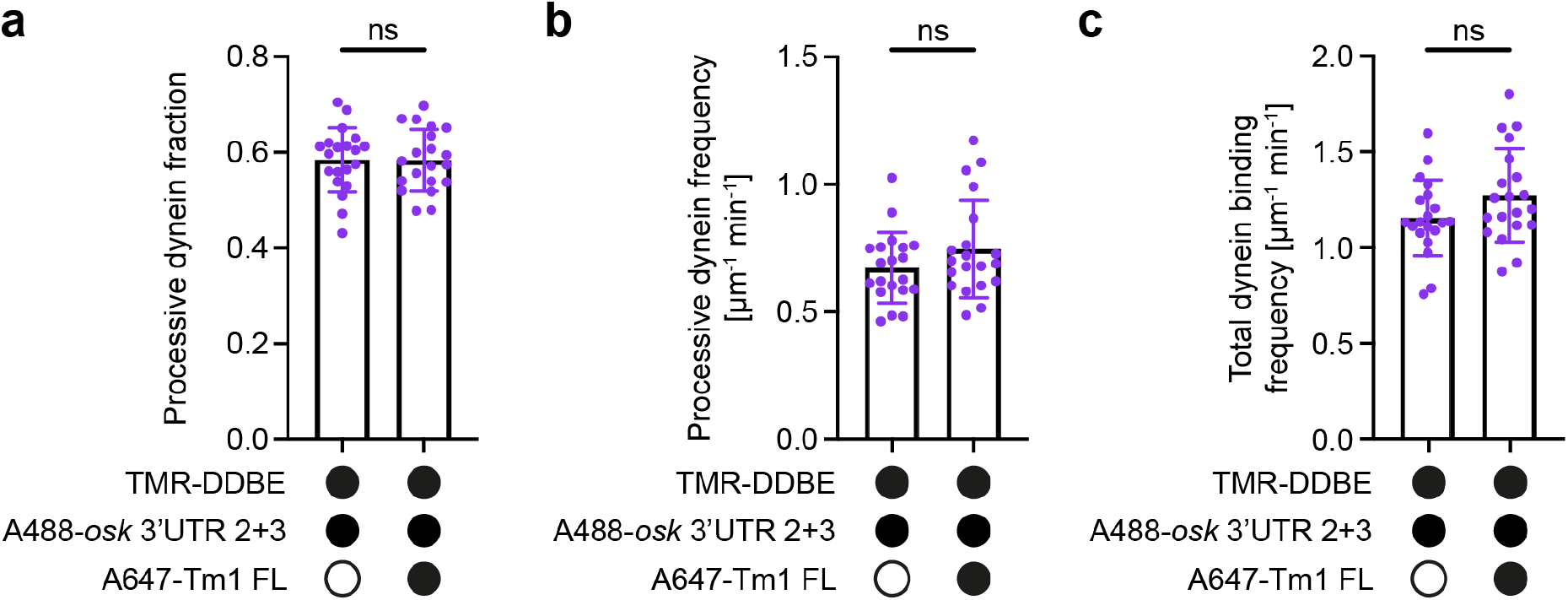
**a-c** Fraction of dynein binding events that underwent processive motility (a) and frequencies of processive dynein movement (b) and dynein binding to microtubules (c) in the presence and absence of Tm1 FL. Dots represent average values for individual microtubules. In each plot, the mean ± SD is shown. Statistical significance was determined by t-test. ns: not significant.

**Figure S10:**
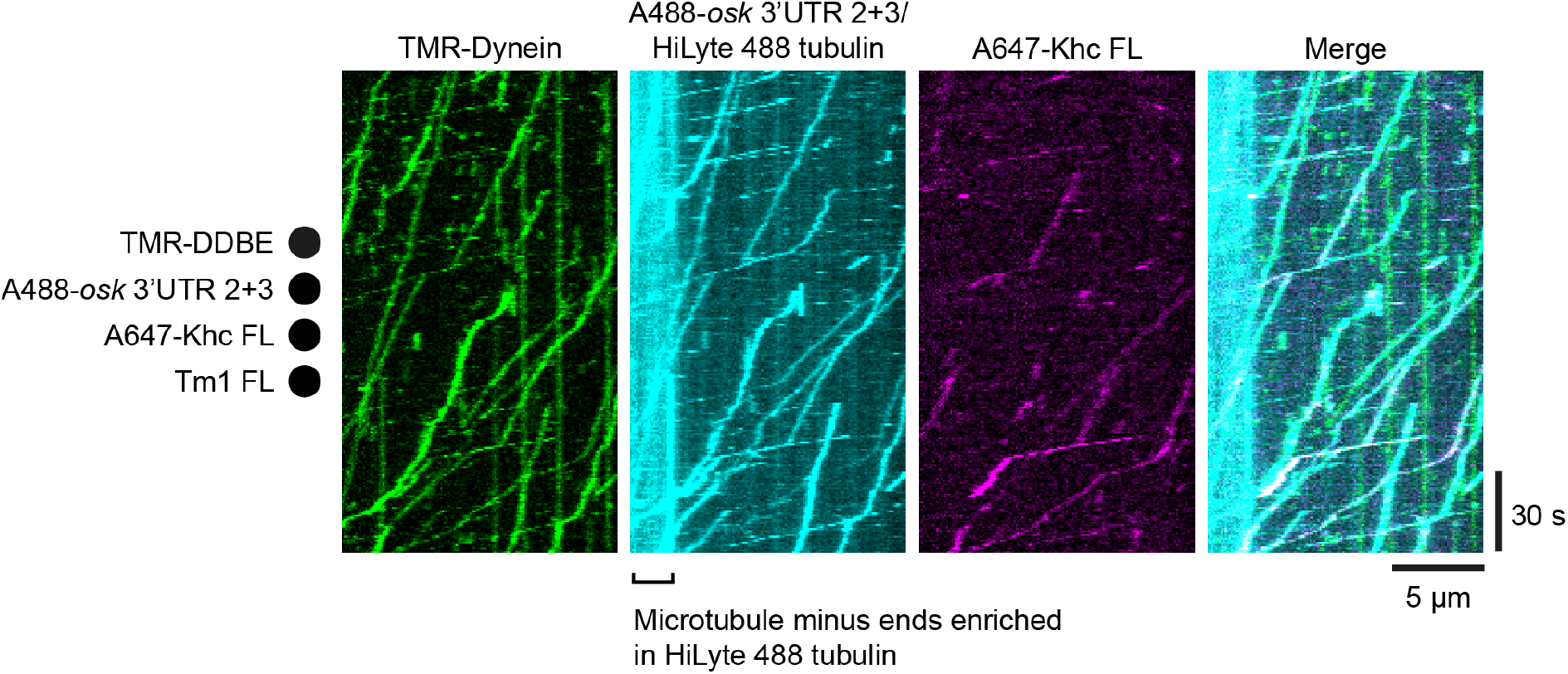
Kymographs showing the Tm1-induced co-translocation of dynein, *osk* 3’UTR 2+3 RNA, and Khc toward microtubule minus ends, which are labeled by enrichment of fluorescent (HiLyte 488) tubulin dimers. Microtubule plus and minus ends are oriented toward the right and left of each kymograph, respectively.

**Figure S11:**
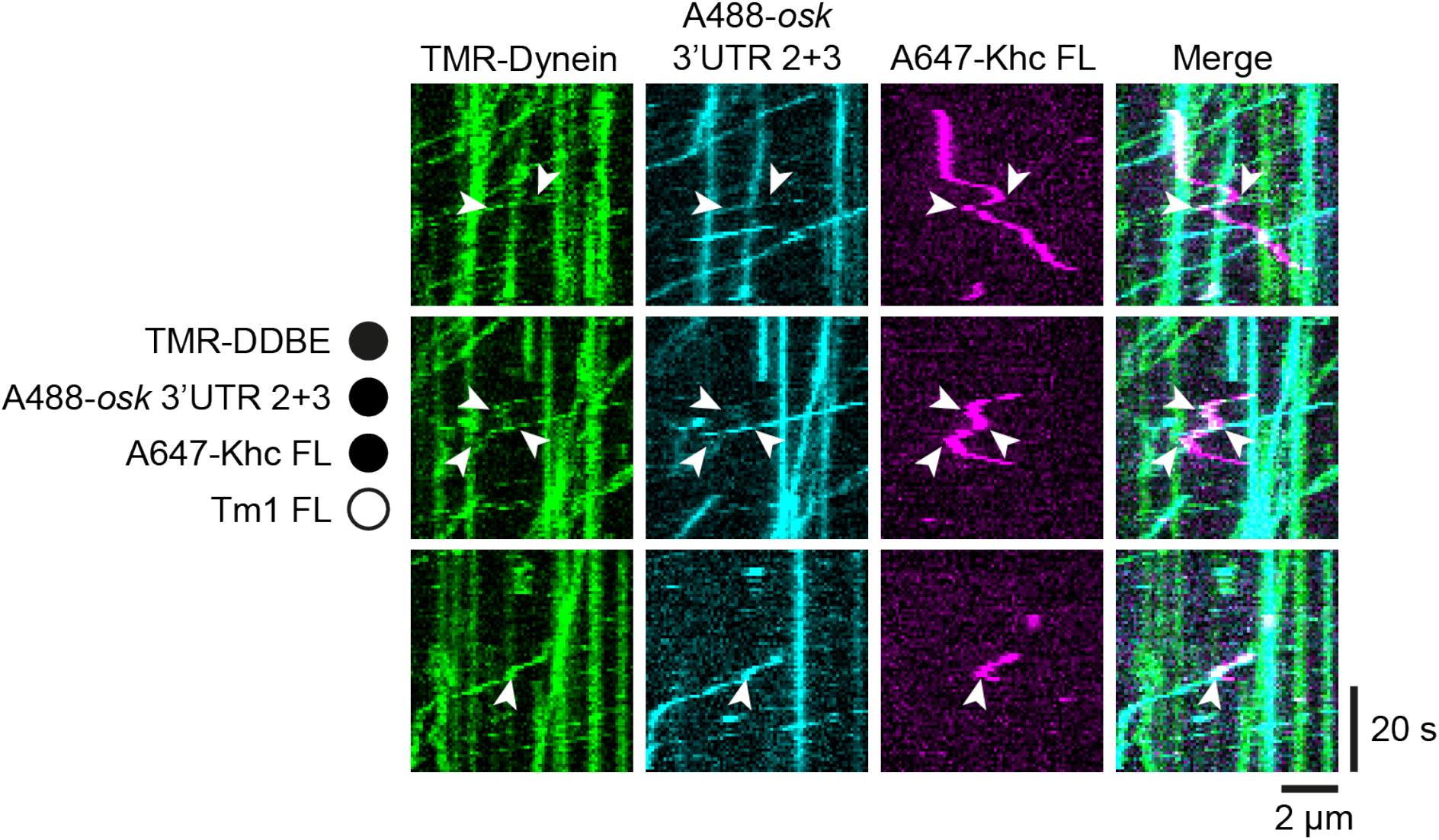
Kymographs showing instances of bidirectional Khc movements in the absence of Tm1. Directional switches of Khc (white arrowheads) correlate with DDBE-*osk* 3’UTR 2+3 association and dissociation events. Microtubule plus and minus ends are oriented toward the right and left of each kymograph, respectively.

**Figure S12:**
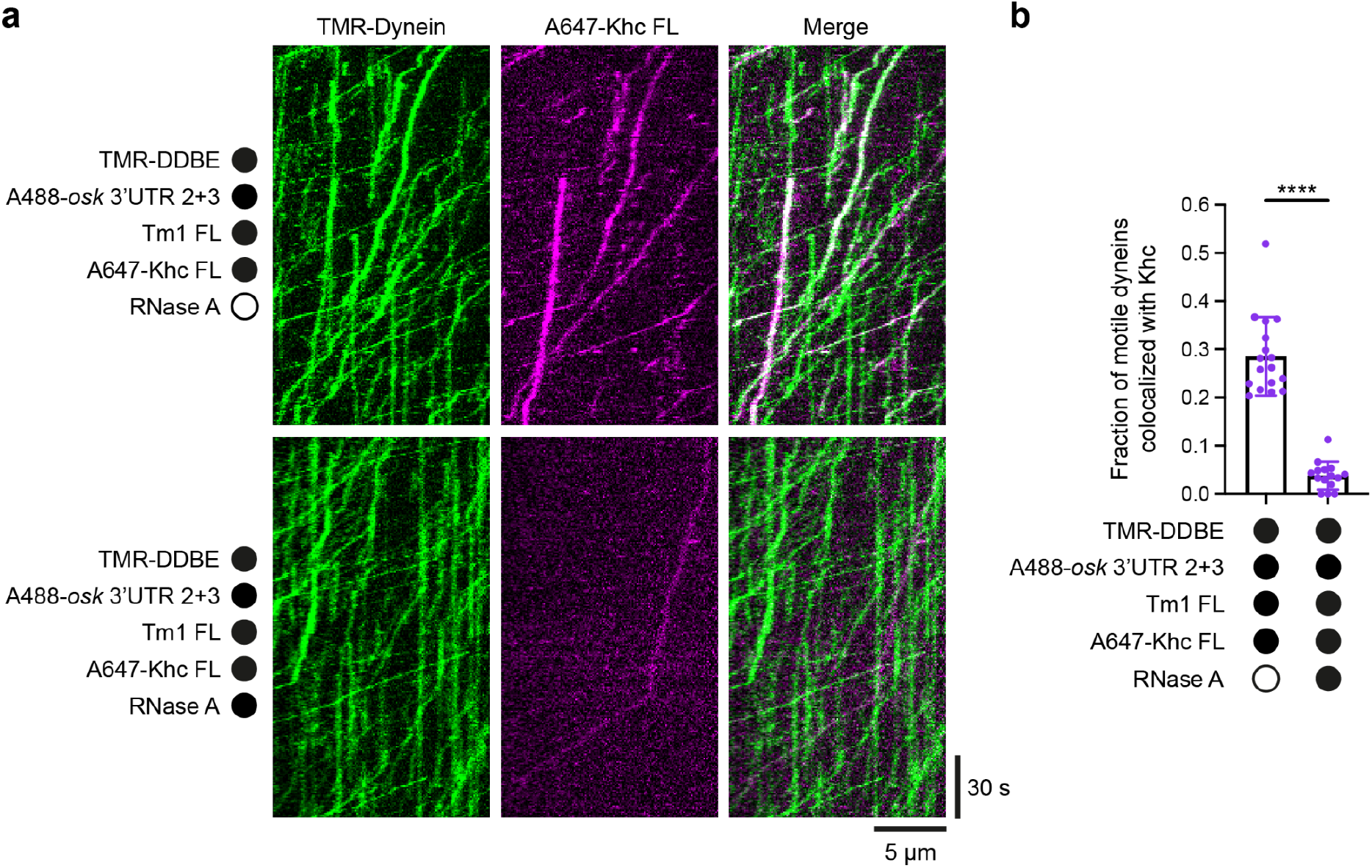
**a** Kymographs showing reduced association of Khc with motile DDBE-*osk* 3’UTR 2+3 RNPs in the presence of Tm1 after treatment with RNase A. Microtubule plus and minus ends are oriented toward the right and left of each kymograph, respectively. **b** Fraction of moving DDBE-*osk* 3’UTR 2+3 RNPs that colocalized with Khc signal with and without RNase A treatment in the presence of Tm1. Dots represent average values of all motility on individual microtubules. Mean ± SD is shown. Statistical significance was determined by Welch’s t-test. ****: p<0.0001.

**Table S1.**
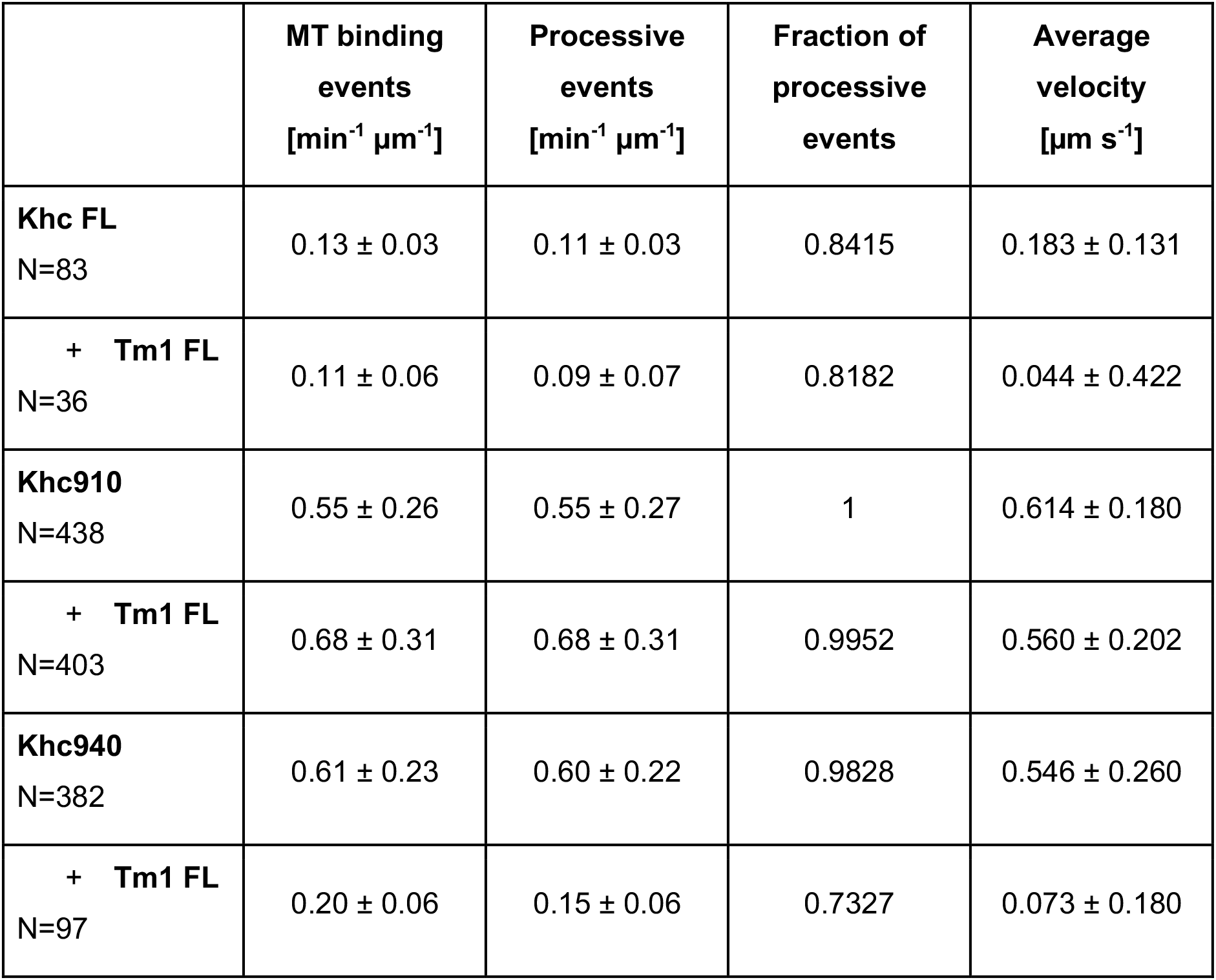
Motility parameters of Khc FL, Khc910 and Khc940

|  | **MT binding**<br>**events**<br>**[min<sup>-1</sup> μm<sup>-1</sup>]** | **Processive**<br>**events**<br>**[min<sup>-1</sup> μm<sup>-1</sup>]** | **Fraction of**<br>**processive**<br>**events** | **Average**<br>**velocity**<br>**[μm s<sup>-1</sup>]** |
| --- | --- | --- | --- | --- |
| **Khc FL**<br>N=83 | 0.13 ± 0.03 | 0.11 ± 0.03 | 0.8415 | 0.183 ± 0.131 |
| + **Tm1 FL**<br>N=36 | 0.11 ± 0.06 | 0.09 ± 0.07 | 0.8182 | 0.044 ± 0.422 |
| **Khc910**<br>N=438 | 0.55 ± 0.26 | 0.55 ± 0.27 | 1 | 0.614 ± 0.180 |
| + **Tm1 FL**<br>N=403 | 0.68 ± 0.31 | 0.68 ± 0.31 | 0.9952 | 0.560 ± 0.202 |
| **Khc940**<br>N=382 | 0.61 ± 0.23 | 0.60 ± 0.22 | 0.9828 | 0.546 ± 0.260 |
| + **Tm1 FL**<br>N=97 | 0.20 ± 0.06 | 0.15 ± 0.06 | 0.7327 | 0.073 ± 0.180 |

**Table S2.** Motility parameters of KhcΔAMB compared to Khc FL and Khc910

|  | **MT binding**<br>**events**<br>**[min<sup>-1</sup> μm<sup>-1</sup>]** | **Processive**<br>**events**<br>**[min<sup>-1</sup> μm<sup>-1</sup>]** | **Fraction of**<br>**processive**<br>**events** | **Average**<br>**velocity**<br>**[μm s<sup>-1</sup>]** |
| --- | --- | --- | --- | --- |
| **Khc FL**<br>N=142 | 0.07 ± 0.15 | 0.05 ± 0.02 | 0.7696 | 0.429 ± 0.300 |
| **+ Tm1 FL**<br>N=24 | 0.02 ± 0.01 | 0.01 ± 0.01 | 0.3114 | 0.159 ± 0.188 |
| **Khc910**<br>N=384 | 0.47 ± 0.10 | 0.47 ± 0.10 | 0.8180 | 0.630 ± 0.169 |
| **+ Tm1 FL**<br>N=595 | 0.73 ± 0.11 | 0.72 ± 0.11 | 0.8955 | 0.649 ± 0.164 |
| **KhcΔAMB**<br>N=65 | 0.06 ± 0.02 | 0.05 ± 0.02 | 0.9929 | 0.507 ± 0.325 |
| **+ Tm1 FL**<br>N=293 | 0.12 ± 0.05 | 0.10 ± 0.04 | 0.9947 | 0.548 ± 0.266 |

**Table S3.** Motility parameters of Khc FL in presence of different Tm1 constructs.

|  | **MT binding**<br>**events**<br>**[min<sup>-1</sup> μm<sup>-1</sup>]** | **Processive**<br>**events**<br>**[min<sup>-1</sup> μm<sup>-1</sup>]** | **Fraction of**<br>**processive**<br>**events** | **Average**<br>**velocity**<br>**[μm s<sup>-1</sup>]** |
| --- | --- | --- | --- | --- |
| **Khc FL**<br>N=335 | 0.23 ± 0.08 | 0.16 ± 0.06 | 0.6847 | 0.32 ± 0.245 |
| **+ Tm1 FL**<br>N=149 | 0.21 ± 0.07 | 0.09 ± 0.04 | 0.4449 | 0.229 ± 0.280 |
| **+ Tm1 1-**<br>**335**<br>N=170 | 0.13 ± 0.03 | 0.06 ± 0.04 | 0.4376 | 0.125 ± 0.165 |
| **Khc FL**<br>N=84 | 0.07 ± 0.02 | 0.06 ± 0.02 | 0.9524 | 0.171 ± 0.212 |
| **+ Tm1 FL**<br>N=44 | 0.04 ± 0.01 | 0.03 ± 0.01 | 0.7137 | 0.188 ± 0.243 |
| **+ Tm1**<br>**252-334**<br>N=171 | 0.08 ± 0.03 | 0.07 ± 0.03 | 0.8694 | 0.303 ± 0.188 |

**Table S4.** Motility parameters of DDBE-*osk* 3’UTR 2+3 in presence and absence of Tm1 FL (Related to Figure 7 and S9). Mean ± SD are shown.

|  | **Total dynein**<br>**binding**<br>**frequency**<br>**[$\mu\text{m}^{-1} \text{min}^{-1}$]** | **Processive**<br>**dynein**<br>**frequency**<br>**[$\mu\text{m}^{-1} \text{min}^{-1}$]** | **Processive**<br>**dynein**<br>**fraction** | **Average**<br>**dynein**<br>**velocity**<br>**[$\mu\text{m} \text{s}^{-1}$]** | **Average**<br>**dynein run**<br>**length [$\mu\text{m}$]** |
| --- | --- | --- | --- | --- | --- |
| **DDBE-*osk***<br>**3'UTR 2+3**<br>**-Tm1 FL** | 1.16 ± 0.20<br>N=20<br>microtubules<br>(2152<br>complexes) | 0.67 ± 0.14<br>N=20<br>microtubules<br>(1277<br>complexes) | 0.58 ± 0.07<br>N=20<br>microtubules<br>(2152<br>complexes) | 0.292 ±<br>0.308<br>N=1277<br>complexes | 4.26 ± 3.90<br>N=1277<br>complexes |
| **DDBE-*osk***<br>**3'UTR 2+3**<br>**+Tm1 FL** | 1.27 ± 0.24<br>N=20<br>microtubules<br>(2207<br>complexes) | 0.75 ± 0.19<br>N=20<br>microtubules<br>(1295<br>complexes) | 0.58 ± 0.06<br>N=20<br>microtubules<br>(2207<br>complexes) | 0.320 ±<br>0.334<br>N=1295<br>complexes | 4.25 ± 3.66<br>N=1295<br>complexes |
| **DDBE-*osk***<br>**3'UTR 2+3**<br>**+Tm1 FL**<br>**(Tm1-bound**<br>**only)** | N/A | N/A | N/A | 0.304 ±<br>0.310<br>N=211<br>complexes | N/A |

**Table S5.** Motility parameters of DDBE-*osk* 3’UTR 2+3 and Khc in presence and absence of Tm1 FL (Related to Figure 8). Mean ± SD are shown.

| | Frequency of Khc transport events toward plus ends [$\mu\text{m}^{-1} \text{min}^{-1}$] | Frequency of Khc transport events toward minus ends [$\mu\text{m}^{-1} \text{min}^{-1}$] | Average Khc-bound dynein velocity [$\mu\text{m s}^{-1}$] | Average distance traveled by Khc toward minus ends [$\mu\text{m}$] |
| --- | --- | --- | --- | --- |
| **DDBE-*osk* 3'UTR 2+3 +Khc -Tm1 FL** | 0.005 ± 0.007<br>N=52<br>microtubules<br>(29 complexes) | 0.014 ± 0.013<br>N=52<br>microtubules (89 complexes) | 0.236 ± 0.223<br>N=109<br>complexes | 3.32 ± 3.82<br>N=109<br>complexes |
| **DDBE-*osk* 3'UTR 2+3 +Khc +Tm1 FL** | 0.005 ± 0.008<br>N=36<br>microtubules<br>(20 complexes) | 0.179 ± 0.064<br>N=36<br>microtubules<br>(822 complexes) | 0.348 ± 0.281<br>N=913<br>complexes | 4.92 ± 4.79<br>N=913<br>complexes |

**Table S6.** Plasmids

| CONSTRUCTS | RELEVANT INFORMATION | SOURCE |
| --- | --- | --- |
| pFastBacDual-HisSUMO-SNAP-3C-Khc | Amp<sup>r</sup>, p10/Polyhedrin promoter | This study |
| pFastBacDual-HisSUMO-SNAP-3C-Khc910 | Amp<sup>r</sup>, p10/Polyhedrin promoter | This study |
| pFastBacDual-HisSUMO-SNAP-3C-Khc940 | Amp<sup>r</sup>, p10/Polyhedrin promoter | This study |
| pFastBacDual-HisSUMO-SNAP-3C-Khc ΔAMB | Amp<sup>r</sup>, p10/Polyhedrin promoter | This study |
| pFastBacDual-HisSUMO-SNAP-3C-Khc-Tm1 | Amp<sup>r</sup>, p10/Polyhedrin promoter | This study |
| pGEX6.1-Khc 855-975 | Amp<sup>r</sup>, tac promoter | This study |
| pETM11-His6-SUMO-SNAP-Tm1-FL | Kan<sup>r</sup>, T7 promoter, lac operator | This study |
| pETM11-His6-SNAP-SUMO-Tm1 1-335 | Kan<sup>r</sup>, T7 promoter, lac operator | This study |
| pETM11-His6-SUMO-Tm1-FL | Kan<sup>r</sup>, T7 promoter, lac operator | 43 |
| pETM11-His6-SUMO-Tm1 1-335 | Kan<sup>r</sup>, T7 promoter, lac operator | 43 |
| pETM11-His6-SUMO-Tm1 252-334 | Kan<sup>r</sup>, T7 promoter, lac operator | 43 |
| pDyn1 (pAceBac1-ZZ-TEV-SNAPf-Dynein heavy chain - DYNC1H1) | Gen<sup>r</sup>, Polyhedrin promoter | 8 |
| pDyn2 (pIDC-Dynein intermediate chain - DYNC1/2, Dynein light intermediate chain - DYNC1LI2, Tctex - DYNLT1, Robl - DYNLRB1, LC8 - DYNLL1) | Chlor<sup>r</sup>, Polyhedrin promoter | 8 |
| pAceBac1-Egl-TEV-ZZ | Gen<sup>r</sup>, Polyhedrin promoter | 15 |
| pIDC-BicD | Chlor<sup>r</sup>, Polyhedrin promoter | 15 |

## RNA sequences

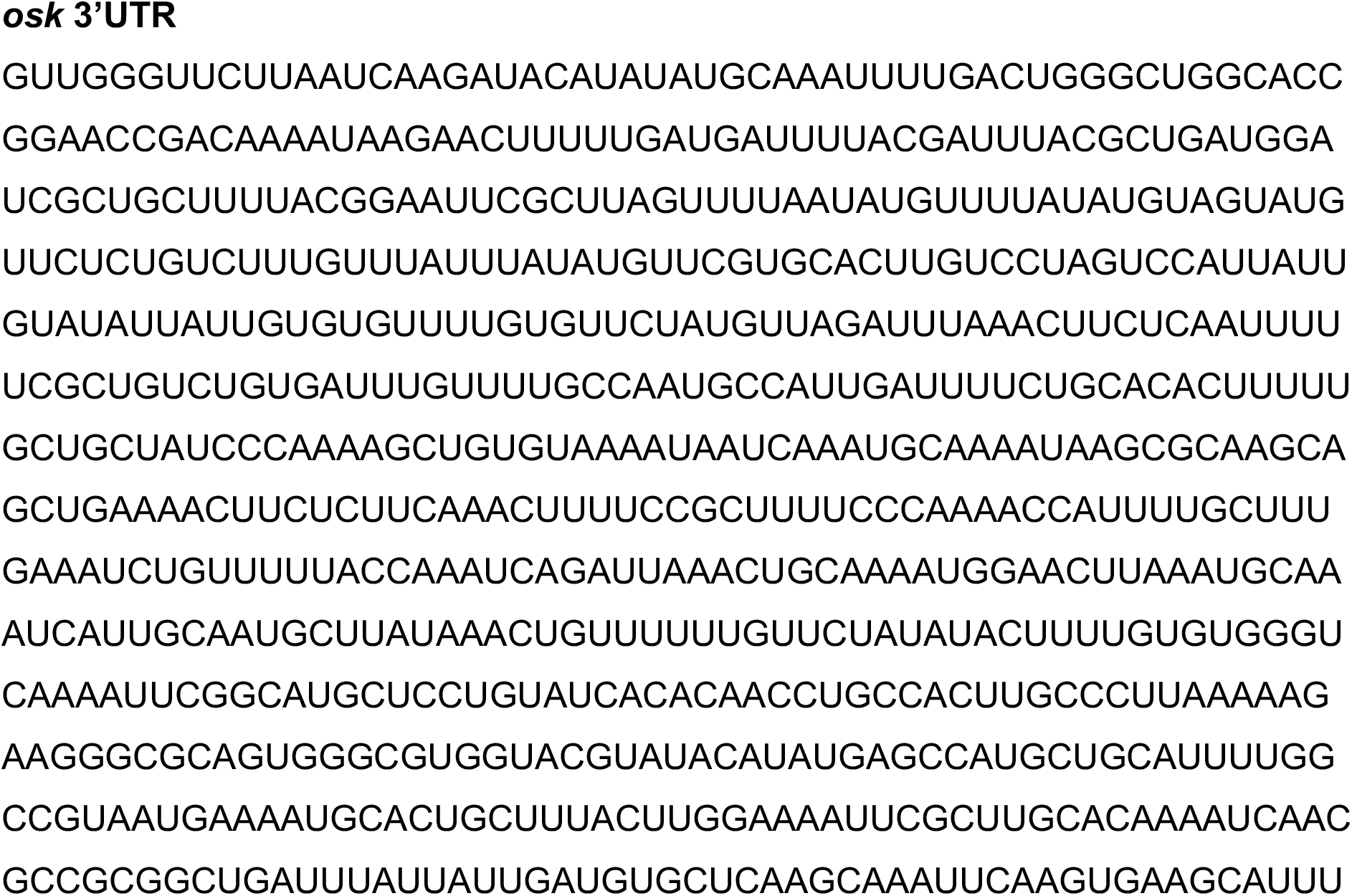
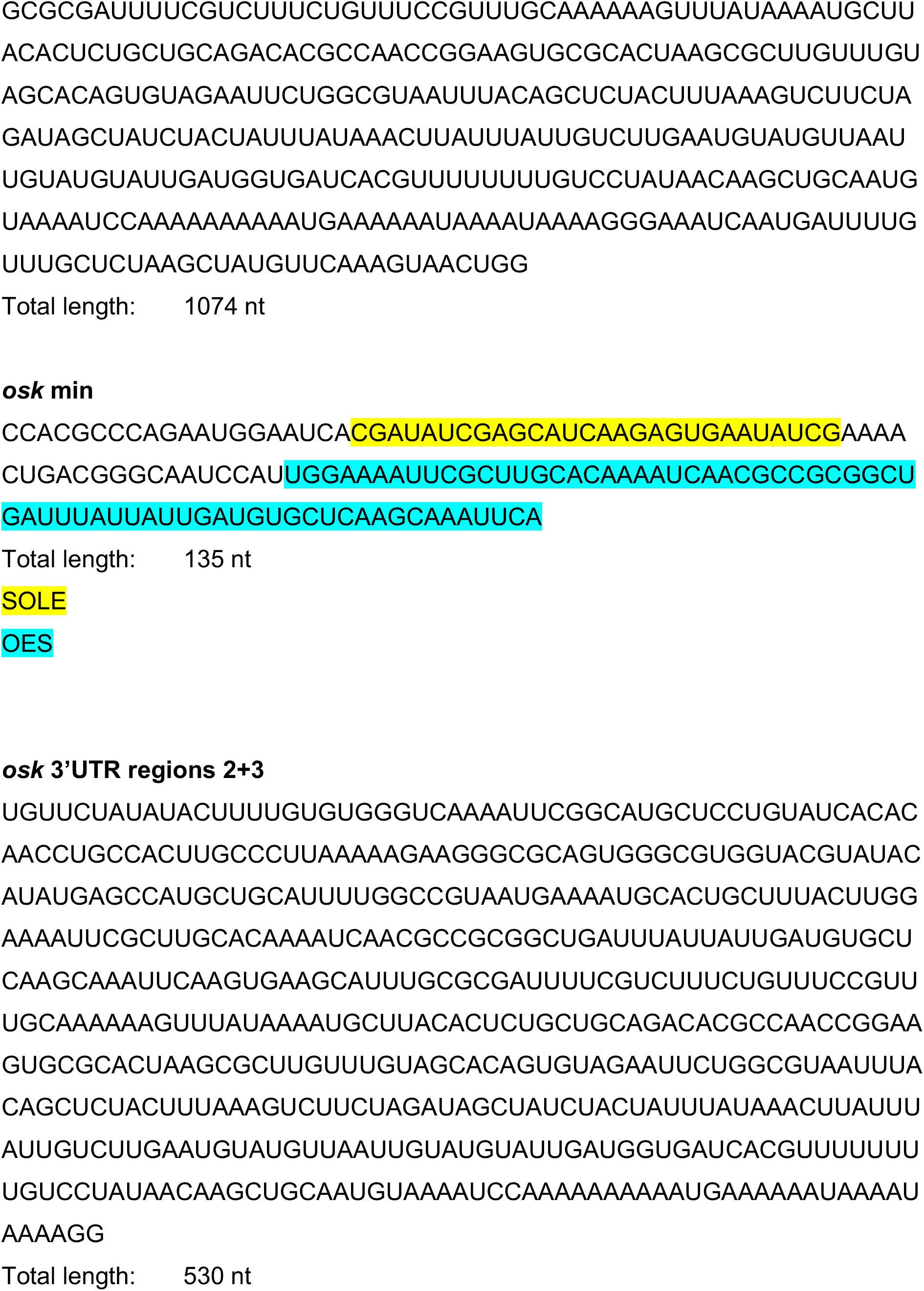

